# Synteny-aware microbial pangenome graphs reveal blueprints of genomic variation

**DOI:** 10.64898/2026.07.03.736256

**Authors:** Alexander Henoch, Metehan Sever, Sarah J. Tucker, Florian Trigodet, Iva Veseli, Tianyi Chang, James O. McInerney, Arda Söylev, Kelle C. Freel, Michael S Rappé, A. Murat Eren

## Abstract

Pangenomics quantifies the conserved and variable gene repertoire among genomes, but popular implementations ignore gene synteny. Graph-based approaches incorporate both gene homology and synteny, but become difficult to interpret due to pervasive rearrangements. Here we present network-pruning and graph-layout algorithms that enable interactive, synteny-aware quantification and visualization of gene conservation and variability. Applied to 29 genomes of the marine genus *Undatipelagibacter* (formerly SAR11 subclade Ia.3.VI), we find that genomic variability forms not a few hypervariable islands against a static backbone but a structured continuum, whose variable regions differ in scale, topology, function, and evolutionary character. Genome variation spans from ancient, specialized regions of hundreds of genes whose propensity to vary is conserved across genera, to single hypervariable genes shaped by epistatic co-selection with partners dispersed genome-wide, and shows that chromosomal context carries evolutionary information synteny-unaware pangenomics cannot capture, and some evolutionary processes act on entire functional subsystems throughout a pangenome.

## Introduction

Microbial genomes are extraordinarily dynamic. Whether through genome streamlining for increased metabolic efficiency (Giovannoni et al. 2014) or genome expansion for increased metabolic versatility (Guieysse and Wuertz 2012), the presence, absence, and organization of genes in genomes can change rapidly through horizontal gene transfer events, integron-mediated cassette shufflings, gene losses, gene duplications, chromosomal rearrangements, and more (Mazel 2006; Koonin and Wolf 2008). Furthermore, individual genes can diversify at different rates among the members of a single population through diversity-generating retroelements (Macadangdang et al. 2022), phase variation (Phillips et al. 2019), or point mutations produced by conventional evolutionary processes. As this constant dynamism enables rapid microbial adaptation to changing environments, understanding the drivers and targets of genome variability promises direct insights into the genetic determinants of fitness with implications that span basic science, medicine, and biotechnology (Eren and Banfield 2024).

One of the most influential observations of comparative genomics has been the prevalence of genomic islands in microbial genomes (Juhas et al. 2009), regions where variation and turnover in gene content typically exceeds genomic averages (Oliveira et al. 2017). Such hypervariable genomic islands are common in host-associated organisms such as *Bacteroides* (Patrick et al. 2010), *Staphylococcus* (Atxaerandio-Landa et al. 2025), and *Rhizobium* (Arashida et al. 2022), as well as free-living environmental bacteria such as *Magnetospirillum* (Ullrich et al. 2005), *Prochlorococcus* (Coleman et al. 2006) and *Pelagibacter* (Wilhelm et al. 2007). Hypervariable genomic islands generally appear to encode context-specific adaptive traits for a given lifestyle. For instance, in host-associated systems they encode antibiotic resistance genes (Schmidt and Hensel 2004) and means to evade host immunity (Liu et al. 2025), and in the environment they encode defense against viruses (Makarova et al. 2011) and nutrient acquisition genes (Hackl et al. 2023).

The lifestyle-critical and niche-specific roles of hypervariable genomic islands suggest that resolving genomic variability and dynamism is necessary for understanding how microbes respond to ecological and evolutionary pressures, and which pressures are most actively (re)shaping their genomes. Yet, the conceptual framework around hypervariability typically assumes that variation in microbial genomes occurs in a bimodal fashion, where stable core regions are interrupted by regions of high variability. This dichotomy yields qualitative rather than quantitative descriptions of overall variability in genomes, and overlooks subtle variation that can neither be classified into stable genomic backbones nor into prominent islands of hypervariability. Plethora of evidence suggests that considering genomic variability within a syntenic framework may provide deeper insights into the presence or absence of organizational principles associated with variability, epistatic interactions between genes and gene neighborhoods that promote or accommodate variation, and their association with selective processes that operate on a lineage (Kang et al. 2014; Bobay and Ochman 2017; Kuronen et al. 2024). However, preserving the syntenic context and making it accessible for downstream exploratory analyses has been a persistent challenge for contemporary comparative genomics.

Pangenomics offers a powerful framework to quantify genomic variation across closely related organisms (Medini et al. 2005), and has become an essential tool in comparative genomics. Standard tools for microbial pangenomics partition genes in a given set of genomes into de novo gene families based on amino acid sequence similarities (Page et al. 2015; Ding et al. 2018; Delmont and Eren 2018). This practical approach yields easy-to-interpret visualizations of gene content variation (see (Delmont and Eren 2018) or (Pena-Fernández et al. 2024) for examples), but it either completely discards the chromosomal context of genes or provides severely limited access to it (Reveillaud et al. 2019), making it difficult to understand how sequence-based gene families relate to their synteny across the genomes represented in a pangenome. One solution for incorporating gene synteny as well as gene homology across genomes lies in representations of pangenomes as gene-centric graph structures (Computational Pan-Genomics Consortium 2018; Sherman and Salzberg 2020), an extension of sequence-centric pangenome graphs that focus on nucleotide-level variation (Paten et al. 2017; Garrison et al. 2018; Li et al. 2020). Several tools implement pangenome graphs that successfully preserve the arrangement of genes across chromosomes (Gautreau et al. 2020; Harling-Lee et al. 2022; Noll et al. 2023; Li et al. 2024), and generic network visualization tools, such as Cytoscape (Shannon et al. 2003), Gephi (Bastian et al. 2009), or Bandage (Wick et al. 2015), offer convenient solutions to display them. However, representing the genomic variability captured in microbial pangenomes poses additional challenges. First, multi-copy genes and genomic rearrangements create complex graph topologies. Second, the use of standard graph layout algorithms that are designed for generic networks do not preserve genomic architecture because they fail to capture the linear nature of chromosomal sequences. Third, the lack of dedicated interactive interfaces limit the exploration of pangenome architectures with the ability to link functions to genes and operons and describe them in genomic regions in which they occur.

Here, we present an open-source software solution to construct, visualize, edit, quantify, and summarize synteny-aware gene-centric pangenome graphs, and apply it to study the genomic variation in *Undatipelagibacter* (previously the Ia.3.VI SAR11 subclade), a recently described genus in the family *Pelagibacteraceae* (Freel et al. 2025), a large group of marine microbes that display remarkable levels of genomic heterogeneity throughout the global ocean (Delmont et al. 2019). The synteny-aware characterization of the *Undatipelagibacter* pangenome reveals an architectural blueprint of this clade in which the variation within non-backbone regions spans a continuum of magnitudes and complexity, and new insights into how structural organization of variation relates to *Pelagibacteraceae* microbial ecology and evolution.

## Results and Discussion

### Graph representation of pangenomes reveals an intricate yet quantifiable genomic variability landscape

Our software solution to study pangenome graphs (1) reduces graph complexity by resolving multi-copy gene ambiguities while decomposing conventional gene clusters (GCs) into synteny-aware gene clusters (SynGCs), (2) untangles complex graphs by resolving cyclic structures into directed acyclic graphs (DAGs), (3) uses a novel network layout algorithm to preserve genomic architectures, and (4) includes a dedicated interactive interface that provides access to functional and metabolic insights into the genes encoded within any given set of paths on the graph (Materials and Methods).

To investigate how conventional pangenomes compare to pangenome graphs in practice, we applied both strategies to compute pangenomes for 29 *Undatipelagibacter* isolate genomes (Grote et al. 2012; Freel et al. 2025). The global prevalence and biogeochemical importance of this surface ocean bacterium, combined with previous reports of fine-scale genomic variation and metabolic adaptations among these 29 co-located isolates from Kāneʻohe Bay, Hawaiʻi (Tucker et al. 2025; Freel et al. 2025), make *Undatipelagibacter* an ideal model for this comparison. Genomes in our dataset had an average length of 1.46 Mbp (1.43 Mbp to 1.54 Mbp), an average GC content of 29.3%, and encoded 1,519 genes on average (1,470 to 1,626) (Supplementary Table 1). The genomes were closely related but not identical: the average nucleotide identity (ANI) between all pairs was 95.1% (Supplementary Table 1). The most similar genome pair in our dataset, HIMB1493 and HIMB1770, had an ANI of 99.99%, while the most distant, HIMB1506 and HIMB1685, had an ANI of 93.4% (Supplementary Table 1).

The organization of genes differed substantially between the conventional *Undatipelagibacter* pangenome and the *Undatipelagibacter* pangenome graph (Figure 1). Despite these structural differences, the overall quantitative summaries produced by the two approaches were broadly similar: for the 43,961 gene calls across the 29 *Undatipelagibacter* genomes, we identified 3,604 GCs and 3,872 SynGCs (Supplementary Table 2), the latter reflecting the higher resolving power of the synteny-aware framework (Materials and Methods). However, because conventional pangenomes lack synteny information, core GCs share identical presence/absence patterns and therefore grouped together, obscuring the spatial relationships between genes (Figure 1). In contrast, the pangenome graph preserved genomic architecture, immediately revealing a consistent pattern of variable regions alternating with backbone stretches of different lengths (Figure 1). The network layout algorithm rendered the genomic landscape captured in the graph more accessible when compared to PPanGGOLiN, a state-of-the-art tool that generates pangenome graphs (Gautreau et al. 2020), which produced a tangled graph from the same set of genomes (Supplementary Figure 1). Of all GCs in the conventional pangenome, 33.6% (1,212) were core GCs, 28.2% (1,016) were accessory GCs, and 38.2% (1,376) were singleton GCs. In contrast, of all SynGCs in the graph, 30.6% (1,186) were core SynGCs, 22.1% (8550 were accessory SynGCs, 9.0% (350) were rearranged SynGCs, 2.8% (107) were duplicated SynGCs, and 35.5% (1,374) were singleton SynGCs (Figure 1, Supplementary Table 2). Critically, the pangenome graph split 189 GCs into 457 SynGCs, exposing the duplicated and rearranged nature of these genes across genomes and substantially improving the resolution of the accessory genomic landscape (Figure 1). Overall, the pangenome graph revealed that only about one third of the *Undatipelagibacter* gene families occurred in stable backbone genomic regions in identical orders across all genomes without interruption, a pattern that was impossible to discern in the conventional pangenome.

**Figure 1:**
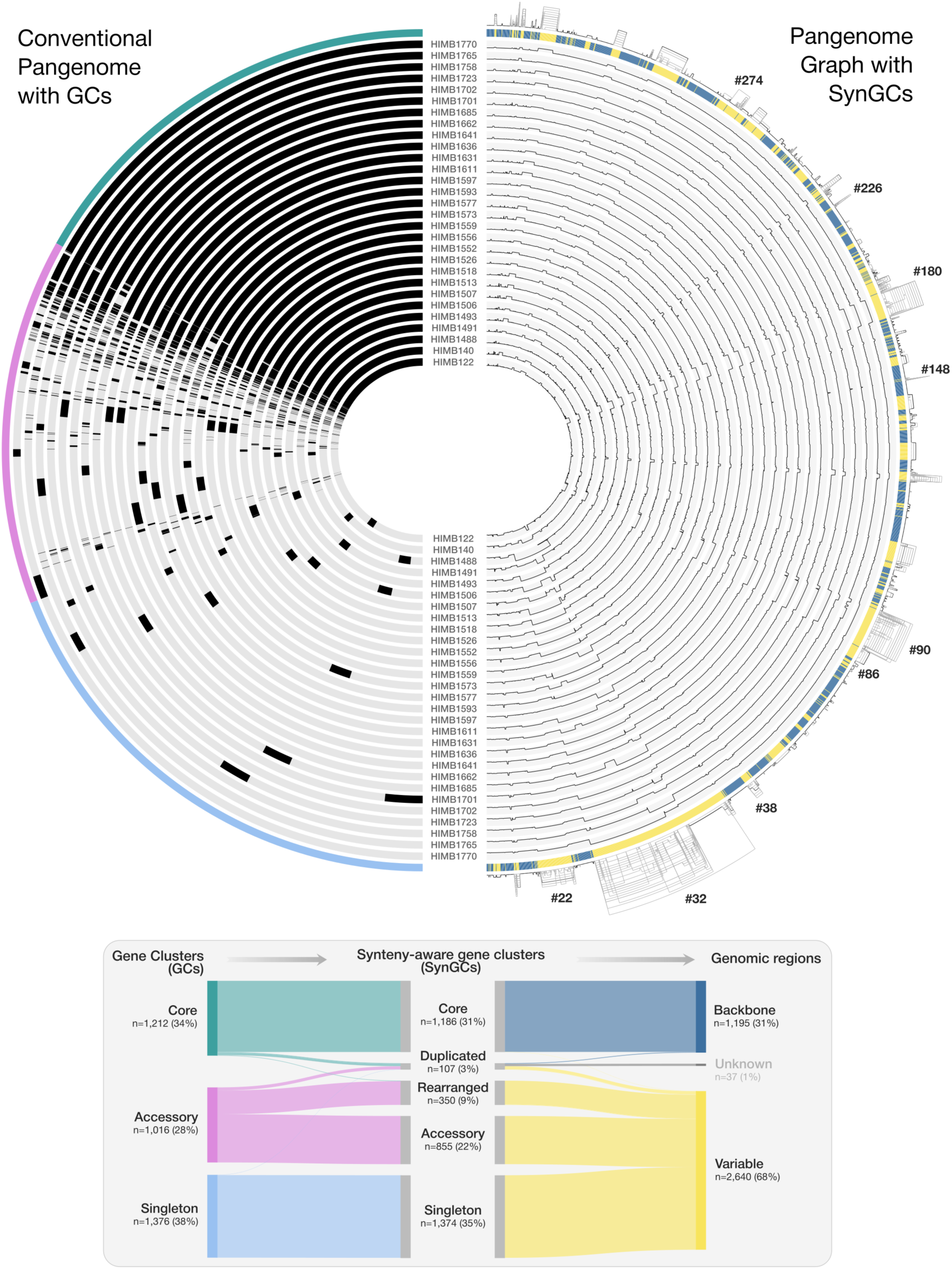
Comparing the *Undatipelagibacter* conventional pangenome and the pangenome graph. The top-left panel shows the conventional pangenome of *Undatipelagibacter* where core, accessory, and singleton gene clusters are marked across 29 genomes in green, pink, and blue colors, respectively. The top-right panel shows the *Undatipelagibacter* pangenome graph with stable backbone regions across 29 genomes marked in blue and variable regions marked in yellow. Variable regions that are mentioned throughout the text are identified by unique variable region IDs. The bottom panel shows a Sankey plot depicting how gene clusters in the conventional pangenome (GCs) relate to synteny-aware gene clusters (SynGCs) and how SynGCs relate to variable and backbone regions of the genome.

### Individual variable regions are functionally specialized and distinct from the genomic backbone

To further investigate the differences between backbone regions (BRs) and variable regions (VRs), we next focused on the functional annotation of SynGCs by grouping all functions described in the NCBI’s Clusters of Orthologous Groups (COGs) database (Galperin et al. 2025) into five broad categories (Supplementary Table 3). While BRs were enriched in functions related to Metabolism & Energy Production and Genetic Information Processing, VRs were enriched with genes that encoded functions related to Cellular Structure & Organization, or those that lacked any annotation (Supplementary Figure 2). These observations are consistent with previous studies which demonstrated that housekeeping functions are enriched in the core genome while environment-facing functions dominate the accessory gene pool in *Pelagibacterales* (Grote et al. 2012) and other marine microbes, including *Prochlorococcus* (Delmont and Eren 2018), and *Alteromonas* (López-Pérez and Rodriguez-Valera 2016).

Taking advantage of the pangenome graph that partitioned the overall genomic variability into distinct VRs that were flanked by backbone genes (Figure 1), we investigated whether individual variable regions were composed of similar functions or differed in their functional profile, and whether the functional composition of VRs consistently differed from the functional composition of the backbone. For this analysis, we excluded 249 regions that showed complete agreement among the contributing genomes, either as conserved backbone regions (171 BRs) or as insertions/deletions (78 INDELs).

The vast majority of VRs were composed of one or two functional categories (Figure 2, Supplementary Table 4). We divided VRs into eight clusters based on their functional composition (Figure 2). The largest cluster (C6), which contained 33 out of 92 remaining VRs, was dominated by genes with unknown functions (Figure 2), which is a well-documented feature of genomic islands in general (Hsiao et al. 2005). The majority of the remaining clusters (C4, C5, C7 and C8) were primarily dominated by Cellular Structure and Organization, Genetic Information Processing, Metabolism and Energy Production, or Cellular Communication and Defense (Figure 2, Supplementary Table 4). While clusters C1, C2, and C3 visually appeared more mixed, they were nonetheless still dominated by two categories rather than a balanced distribution (Supplementary Table 4). These data suggest a high degree of within-VR functional cohesion, and point to distinct and coherent functional identities across VRs.

**Figure 2:**
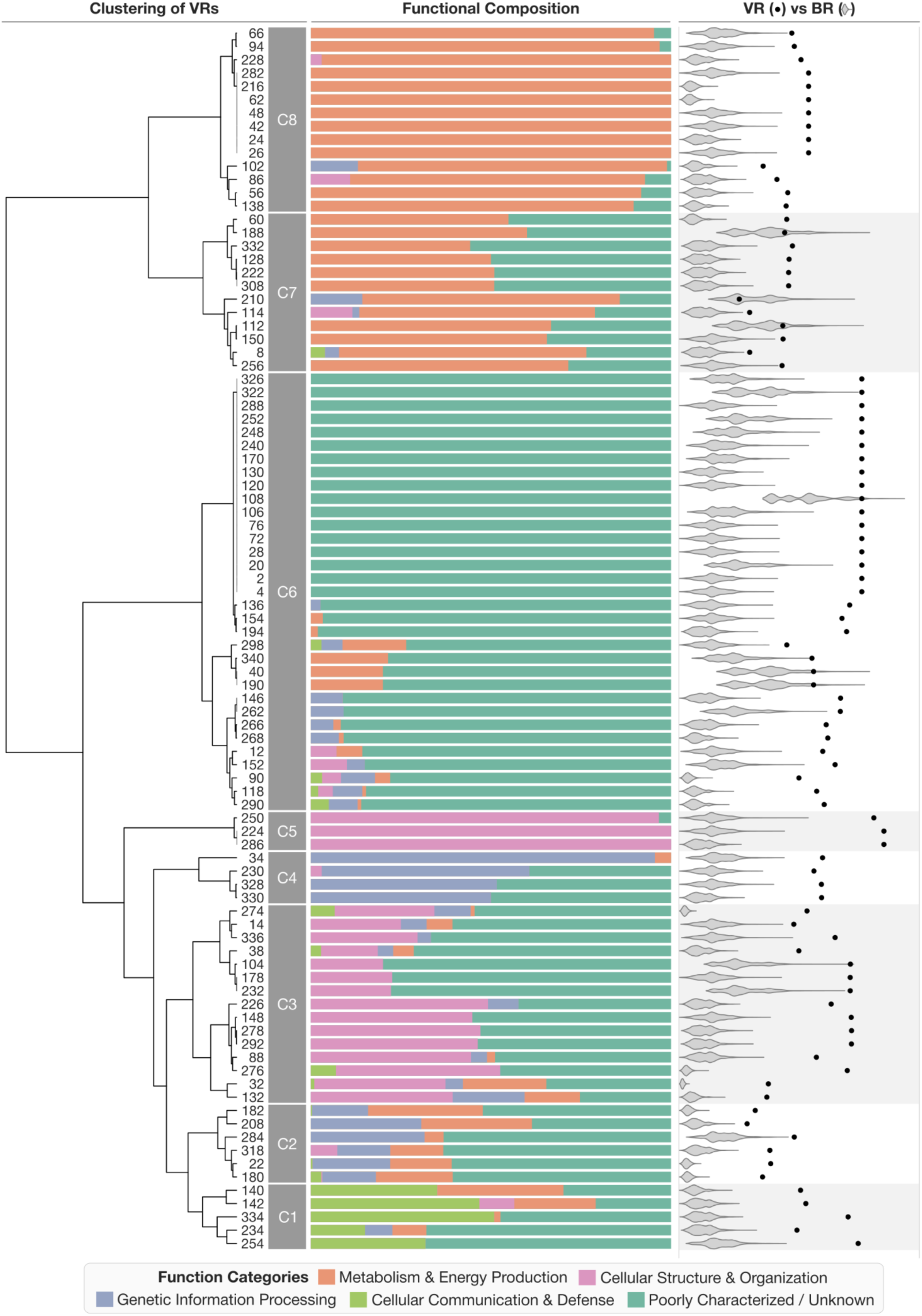
Functional distribution of diverse variable regions. Barplots show the distribution of the five major groups of COG24 functional categories (Metabolism & Energy Production, Cellular Structure & Organization, Genetic Information Processing, Cellular Communication & Defense, and Poorly Characterized or Unknown) within a given variable region (VR), and the dendrogram on the left organizes VRs based on their functional composition. Black dots show the Hellinger Distance between the functional distribution of a given VR and backbone regions (BRs), while the adjacent violin plots show the distribution of the same distance between that VR and 10,000 randomly sampled BRs of equivalent length, providing a null expectation.

To test whether the functional composition we observed in VRs distinguished them from backbone regions, we compared the proportion of functions encoded in each VR to the same number of genes randomly sampled 10,000 times from the backbone. This analysis showed that the functional composition of nearly all VRs was significantly different from that of the backbone (Hellinger distance, p < 1e-5), with the exception of nine that showed overlap with the null distribution (Figure 2, Supplementary Table 4). Together, these findings suggest that the variable regions in the *Undatipelagibacter* pangenome are not genomically diverse in a random sense, but represent functionally specialized and coherent units that are distinct from the genomic backbone.

The functional coherence of individual VRs raised a separate question: do VRs of similar character occupy similar positions across related lineages? To investigate this, we revisited the original description of SAR11 genomic islands by Wilhelm et al. (2007), who described four hypervariable regions in *Pelagibacter* (formerly SAR11 subclade Ia.1.I), another genus of the family *Pelagibacteraceae* (Freel et al. 2025). The authors noted that the most variable region in their analysis, so-called HVR2, was flanked by rRNA genes and was enriched in cell envelope biogenesis, outer membrane transport, and genes of unknown function (Wilhelm et al. 2007). The largest and most functionally distinct VR in our analysis of the *Undatipelagibacter* pangenome, so-called VR #32 (Figure 1), showed a near-identical functional signature to HVR2, and was similarly flanked by rRNA and tRNA genes (Supplementary Table 5, Supplementary Figure 3). This correspondence between two distinct genera in the *Pelagibacteraceae* compelled us to survey whether similarly complex VRs with similar functional characteristics occurred in similar genomic contexts in other genera as well. For this, we constructed pangenome graphs for four additional genera in the *Pelagibacteraceae* and one additional genus in the *Fontibacteraceae* that each had at least three high-quality genomes. We found that each of them indeed carried a VR of similar complexity and functional make-up located close to the DnaA gene and flanked by an rRNA and three tRNAs (Figure 3, Supplementary Figure 3). The overall graph structure of these HVR2-analogues, which were consistently the most variable region within each genus of the *Pelagibacterales*, differed considerably across genera. However, their composition and location remained consistent throughout all genomes in our analyses (Figure 3), as other researchers also observed through analyses of different sets of genomes from the same order (Sadler et al. 2025; Nielsen and Lui 2026).

**Figure 3:**
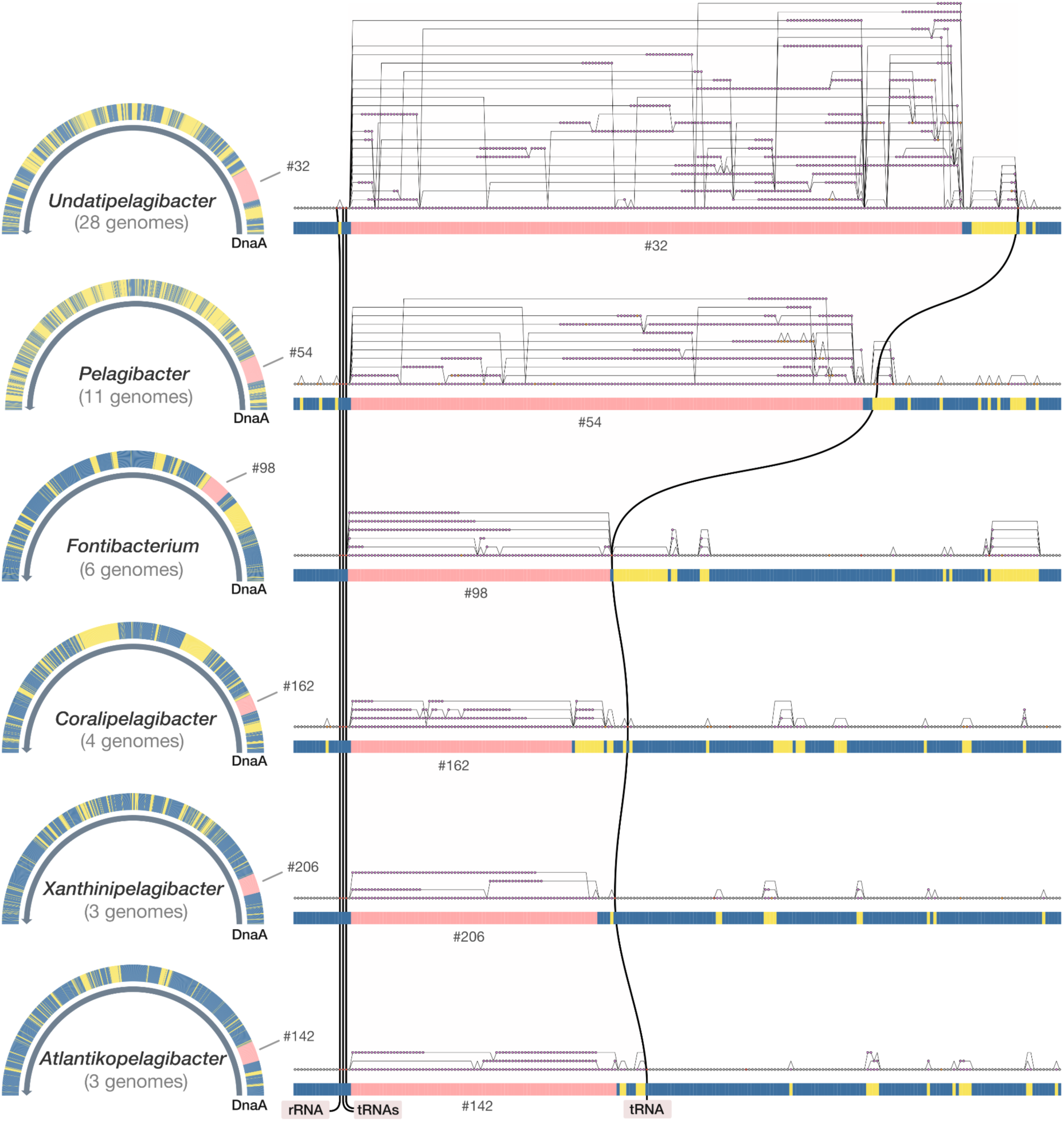
Comparison of the most variable genomic regions within the *Pelagibacterales*. For each genus, the left panel shows the broader context of the variable region (marked in pink) while the right panel offers a detailed representation of the variable region graph structure. The backbone and variable regions are shown in blue and yellow, while the lines that connect the variable regions across genera mark the location of flanking tRNA and rRNA genes. The visualization of region #32 in *Undatipelagibacter* excludes HIMB1488 to simplify the visualization.

What is conserved here is not the sequence itself, which differs dramatically from genome to genome in this locus, but the variability of the locus, which manifests in the same functional context and location across multiple genera (Figure 3). This conservation of the propensity to vary, rather than the conservation of any particular sequence, suggests that this highly variable locus was already part of the ancestral *Pelagibacterales* genome before these genera diverged, and that its capacity for variation has been maintained since then. Identifying such ‘Persistent Variable Regions’ (PVRs) through integrated analyses of phylogenomics and pangenome graphs can reveal which regions of genomic variability are ancestral features that are maintained across deep evolutionary timescales, and which are more recent acquisitions.

The existence of PVRs has implications for theoretical frameworks of pangenome evolution that model accessory genes individually, with their frequencies shaped by gene gain, gene loss, and gene-specific fitness effects (Niehus et al. 2015; McInerney et al. 2017; Brockhurst et al. 2019). However, the conservation of organized variable loci across genera suggests that selection may act not only on individual gene content, but also on the genomic architecture that organizes variability into discrete, ancestral PVRs. Within such regions, a large number of sequence-discrete genes may ultimately correspond to a relatively small number of functional roles.

### Graph-derived metrics quantify genomic variability as a continuum

While the functional characterization of VRs yields insights into their potential role in fitness, it says nothing about how this variation is arranged across genomes. With its detailed description of the VR topology (i.e., how individual genes branch, reconnect, and route different genomes through alternative paths in the graph), the graph representation allows for measuring the architectural properties of VRs independent of gene function. Indeed, the *Undatipelagibacter* pangenome graph revealed distinct topological motifs within each VR (Figure 1), which we sought to quantify.

We defined four metrics to capture different aspects of a given VR (Materials and Methods): (1) ‘complexity’, the number of paths in the VR subgraph needed to explain differing organizational patterns of SynGCs across genomes, (2) ‘expansion’, the largest number of genes any single genome adds to the VR, (3) ‘diversity’, the variance in gene prevalence among SynGCs in the VR, and (4) ‘weight’, the number of genomes in which the VR occurs, as a fraction of the dataset maximum. We further combined these four metrics to implement a single Composite Variability Score (CVS), which we defined as the geometric mean of diversity, expansion, and complexity multiplied by weight, for quantitative descriptions of the relative variability of each region in a given pangenome graph (Figure 4).

**Figure 4:**
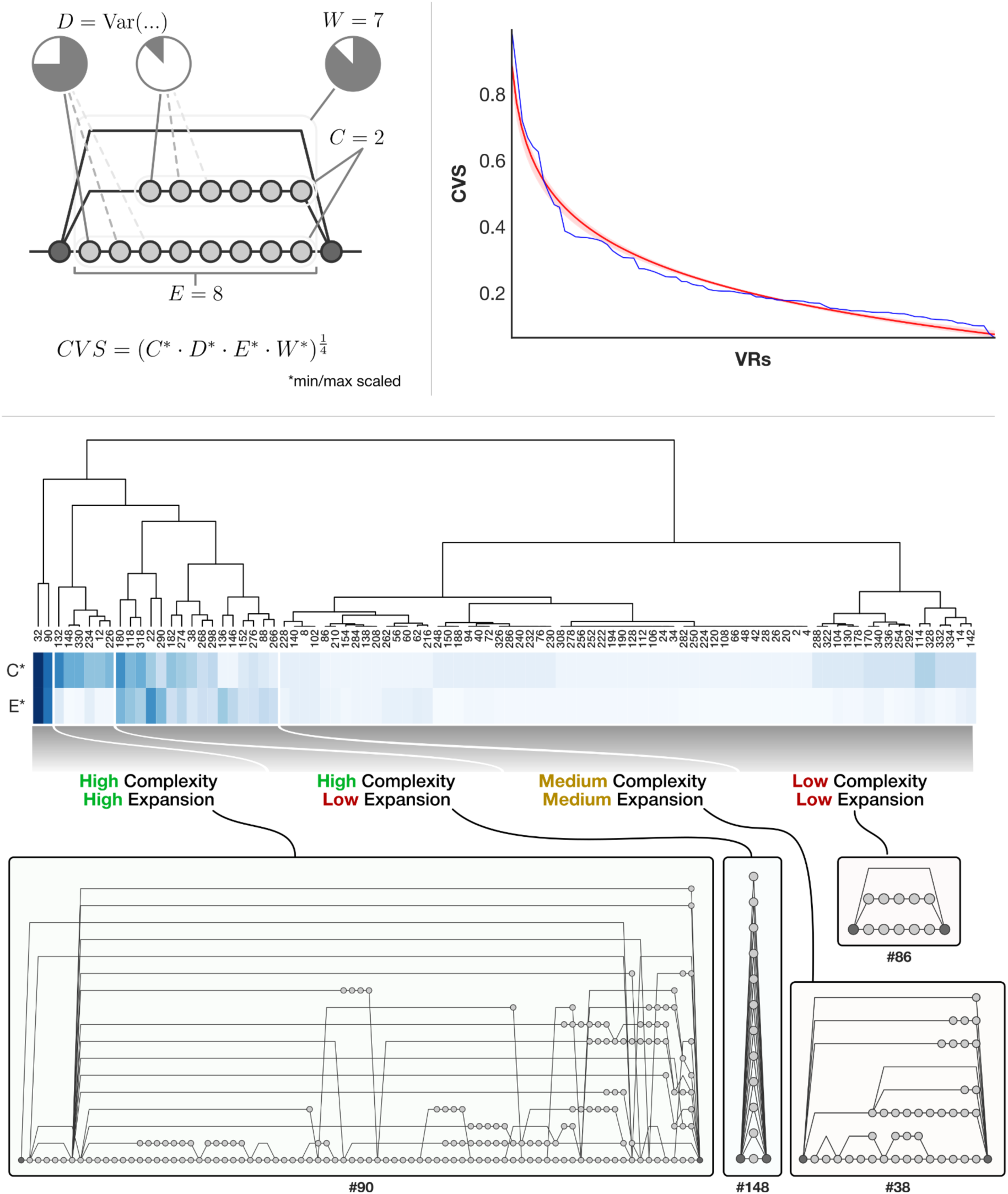
Graph features that contribute to the calculation of the ‘Composite Variability Score’ and the clustering of *Undatipelagibacter* variable regions via pangenome graph metrics. The top-left panel illustrates how complexity (C), expansion (E), diversity (D), and weight (W) are derived from graph features and combined into a Composite Variability Score (CVS) for any given variable region. The top-right panel shows the CVS curve across all variable regions in the *Undatipelagibacter* pangenome graph. The bottom panel shows the hierarchical clustering of VRs based on their complexity and expansion values, in which VRs are grouped into four clusters, and a representative graph region from the *Undatipelagibacter* pangenome is shown below each cluster.

Of the 341 distinct variable and backbone regions in the *Undatipelagibacter* pangenome graph, 92 had a CVS above zero (Figure 4). Four VRs (#32, #90, #22, and #180) stood out with CVS values exceeding 0.5. Each of these VRs was present in all 29 genomes, and they collectively accounted for 58.8% of all SynGCs across variable regions (Supplementary Table 6). The functions encoded in these regions were diverse (Supplementary Table 6). VR #32 was dominated by lipopolysaccharide and O-antigen biosynthesis genes, glycosyltransferases, and NDP-sugar modification enzymes, which are the functional hallmarks of cell envelope biogenesis. VR #90 contained cell surface modification genes, genes for restriction-modification defense systems, and integrases and recombinases. VR #22 included genes for histidine biosynthesis, DNA recombination and repair, and a DNA phosphorothioation defense system. VR #180 included genes for amino acid catabolism and efflux along with envelope stress-response regulation.

In addition to their distinct functional repertoires, the regions with the highest CVS values also exhibited fundamentally different patterns of genomic variation. VR #32 harbored a total of 1,005 SynGCs, representing 40.8% of the total SynGCs across all VRs. With 35 to 88 genes per genome and a gene calls-to-SynGC ratio of 1.7, VR #32 was consistently well represented across genomes, yet most of its gene content belonged to SynGCs unique to individual genomes. In contrast, VR #90, which ranked second by CVS, exhibited a strikingly different pattern of variation as this VR was represented by anywhere from 1 to 65 genes across individual genomes, revealing a much broader range in representation compared to VR #32 across individual members of the *Undatipelagibacter* (Supplementary Table 6). VRs with relatively lower CVS also revealed noteworthy patterns. For instance, VR #274 (CVS = 0.459), which contained genes involved in type IV pilus assembly and secretion, as well as surface adhesion and signal transduction, contained 647 gene calls but only 41 SynGCs, yielding the highest ratio of genes per SynGC (15.8) in the entire dataset by far (Supplementary Table 6). With its very low normalized diversity estimate of 0.096, this VR appeared to be a region where genomes share many of the same genes but simply arrange them differently, making VR #274 a region of structural variation rather than gene content variation, the exact opposite pattern of VR #32. Together, these contrasting examples demonstrate that genomic variability can manifest through fundamentally different forms even within a single pangenome.

The CVS curve showed a continuous decrease without a clear breakpoint that could partition VRs into distinct bins (Figure 4). The CVS distribution was continuous and unimodal (Hartigan’s dip test, p = 0.9), and lacked a statistically supported inflection point in the ranked curve (PELT changepoint analysis). Consistent with these observations, the data were better explained by a single log-normal distribution than by a two-component mixture model (ΔAIC = 13), indicating that genomic variability in *Undatipelagibacter* is more of a continuum rather than a dichotomy between a few hypervariable islands and an otherwise static genome. This continuum is obscured when variable regions are defined using discrete arbitrary quantitative or qualitative thresholds rather than quantified along a continuous scale. This empirical observation resonates with the theoretical prediction that core and accessory genes represent a continuum governed by the proportion of environments in which genes are favored rather than explicitly distinct categories (Niehus et al. 2015; McInerney et al. 2017). However, pangenome graphs reveal that this continuum is not one-dimensional: variable regions differ in structural topology, functional identity, and scale, a level of complexity that frameworks based solely on gene frequency cannot capture.

Despite the continuous distribution of variability, hierarchical clustering of VRs based on their complexity and expansion values revealed at least four broad structural categories with distinct topological properties (Figure 4). VRs in the ‘high complexity / high expansion’ cluster occupied large genomic stretches and contained many unique paths through the graph, representing the most prominent hotspots of genomic variation such as the VRs #32 and #90. VRs in the ‘medium complexity / medium expansion’ and ‘low complexity / low expansion’ clusters showed progressively simpler and more compact graph structures where entire operons were present in some genomes but absent from others. In contrast, VRs in the ‘high complexity / low expansion’ cluster were confined to narrow genomic windows wherein a relatively small number of genes were flanked by backbone regions and yet harbored striking levels of variation across genomes (Figure 4). The relationships between VRs based on their topological properties showed little agreement with their organization based on the functions they encoded (Supplementary Figure 4), suggesting that topology and function likely carry independent information about the evolutionary forces that shape them, at least in *Undatipelagibacter*. Whether this relationship differs among other clades of life warrants further investigation.

### Epistatic co-selection ties the evolution of a chaperone to its clients within and beyond a conserved operon

The *Undatipelagibacter* pangenome graph revealed a strong correlation between expansion and complexity values within VRs (Pearson’s r=0.7, *p* < 1e-13) (Supplementary Table 6), which reflects a general tendency for highly variable regions to also occupy larger genomic footprints. Variable regions with high complexity but low expansion, however, are notable exceptions to this trend as they describe regions with striking levels of sequence divergence across a large number of genomes, yet maintain extremely small footprints that are in some cases as little as a single gene (Figure 4). Region #148 is such a case, where multiple *Skp* genes encoded by highly divergent sequences are flanked by conserved genes (Figure 5).

**Figure 5:**
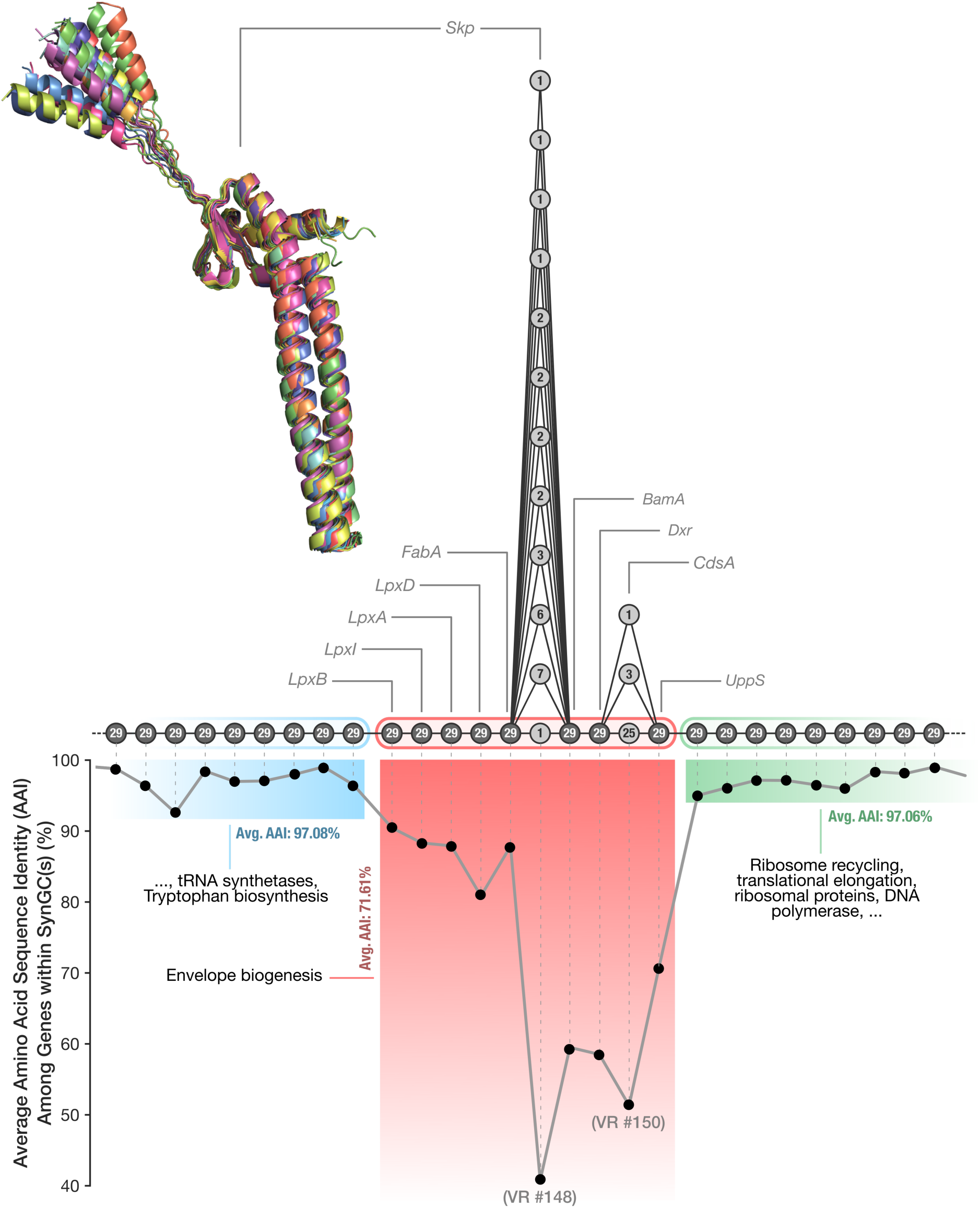
The genomic context and sequence similarity gradient around *Skp.* Each node in the subgraph represents a SynGC, and the numbers in them represent the total number of genomes that contribute a gene to a given SynGC. any of the 29 *Undatipelagibacter* genomes. is represented. The protein structure overlays predicted protein structures we calculated for each *Skp* gene representative from each SynGC. The line graph at the bottom of the figure displays the average amino acid identity (AAI) across gene sequences that occur at the same graph position.

The ten genes surrounding VR #148 form a functionally cohesive envelope biogenesis module that appears to be largely conserved in *Proteobacteria* (Alakavuklar et al. 2023) (Figure 5, Supplementary Table 7). Current research suggests that (1) *LpxA*, *LpxB*, *LpxI*, and *LpxD* are involved in lipid A biosynthesis (Raetz and Whitfield 2002) (with the use of *LpxI* rather than *LpxH*, consistent with previous observations for *Alphaproteobacteria* (Metzger et al. 2012)), (2) *FabA* produces unsaturated fatty acyl chains (lipid A acyl donors) (Parsons and Rock 2013), (3) *Skp* and *BamA* collaborate in the periplasmic chaperoning and outer membrane insertion of beta-barrel proteins (Konovalova et al. 2017), (4) *Dxr* supplies isoprenoid precursors (methylerythritol phosphate pathway) (Frank and Groll 2017), (5) *CdsA* synthesizes the CDP-diacylglycerol branch point of phospholipid biosynthesis (Parsons and Rock 2013), and (6) *UppS* produces the undecaprenyl pyrophosphate lipid carrier, essential for peptidoglycan and LPS assembly (Bouhss et al. 2008). The functional cohesion of genes immediately surrounding VR #148 is reinforced by the observation that the flanking regions encode entirely different functions. Immediately upstream of this region appears a tryptophan biosynthesis operon (*TrpEGDC*) followed by *LexA* and glutamyl-tRNA synthetase, while genes immediately downstream are related to translation and include a ribosome recycling factor, elongation factor Ts, ribosomal protein S2, and DNA polymerase III. These data suggest that the envelope biogenesis module containing the highly variable *Skp* gene resembles an operon flanked by others that serve different purposes (Figure 5).

*Skp* was the only gene in this operon that split into 12 sequence clusters across genomes, whereas nearly all of the other genes maintained sufficient sequence similarity to remain grouped into single clusters. This observation raises an interesting question: does the sharp peak of sequence divergence reflect a true boundary between a single highly variable gene in an otherwise stable operon, or does it represent the point where gradual sequence divergence across the operon finally exceeds the clustering threshold? To address this question, we examined the nature of within-position average amino acid identity (AAI) scores across all genes in a larger genomic context.

Average AAI values revealed a much less uniform variability landscape compared to what is evident by the partitioning of genes into sequence clusters. There was an abrupt, operon-level change in the level of sequence similarity across SynGCs: genes encoded upstream and downstream of the envelope biogenesis operon had an average AAI of 97%, with a minimum average AAI value of 92% (Figure 5, Supplementary Table 7). In contrast, the envelope biogenesis operon had an average AAI of 71%, with a minimum average AAI value of 40% (Figure 5, Supplementary Table 7). However, unlike the relatively stable AAI values in the surrounding regions, the identity scores within the envelope biogenesis operon were not uniform. The relatively high AAI values at the start and end of the operon progressively declined towards its center, forming a sequence similarity gradient that reached its lowest around 40% at the *Skp* gene. Despite the very low sequence similarity, AlphaFold2-predicted protein structures for representatives of all twelve *Skp* SynGCs were highly similar (Figure 5), shown by an average TM-score of 0.9, despite subtle structural differences (Supplementary Figure 5). This observation raises two related questions: What maintains such extensive variation in *Skp* gene sequences when the resulting structures are so similar, and what mechanism produces the gradient in sequence divergence in genes surrounding *Skp*?

The variation in *Skp* itself may be explained by diversifying selection on cell surface structures exerted by phages or grazers that target *Pelagibacterales*, given the way *Skp* sequesters a diverse set of OMPs within an expandable cavity (Schiffrin et al. 2016). Because even slight structural variants may require substantial compensatory changes in the underlying amino acid sequence (Storz 2018), the high sequence divergence among *Skp* genes may reflect structural fine-tuning of this chaperone to match the variation in its clients. But this likely cause behind the high sequence variation among *Skp* genes does not explain the sequence similarity gradient that descends from over 97% AAI in the flanking genes to below 40% at *Skp* (Figure 5). At least two non-mutually exclusive processes could account for this pattern.

The first is staggered homologous recombination. In any single genome, the import of a divergent DNA tract will create a discrete boundary (i.e., high identity in the unrecombined flanks and an abrupt drop across the introgressed segment). But, as recombination breakpoints can fall at different positions in different lineages, averaging many such events across a single population can, in theory, ‘smear’ discrete events into a continuous identity valley, almost a population-scale echo of the position-dependent boundaries that recombination produces (Retchless and Lawrence 2007). We found various lines of evidence in our data that were consistent with this narrative. The distribution of pairwise amino-acid identities at individual operon genes was frequently bimodal, which can be interpreted as evidence for a high-identity ‘native’ mode and a low-identity ‘divergent’ mode. This is an expected outcome when native and introgressed alleles coexist in a population (Birzu et al. 2025), which was absent from the conserved flanking genes (Supplementary Figure 6). Both synonymous and nonsynonymous divergence were elevated across the operon, in agreement with the general expectation when entire DNA tracts move rather than individual sites mutate (Didelot and Wilson 2015). Because the synonymous sites approached saturation, they were not as informative about the precise depth of import, but when we tested the contiguous locus directly with the pairwise homoplasy index (Bruen et al. 2006), the null hypothesis of clonal evolution was rejected (PHI test, p = 1e-4) as phylogenetic incompatibility between polymorphic sites increased with the distance separating them (Supplementary Figure 7). Importantly, this recombination signal was not confined to the operon alone: when we tested the operon and the flanking genes separately under the same procedure (9,999 permutations each), both rejected clonality (PHI; p = 1e-4). This observation is in line with suggestions of pervasive, genome-wide recombination within *Pelagibacterales* (Vergin et al. 2007; Vos and Didelot 2009), and thus it does not say anything specific about the *Skp*-encoding operon. When we tested whether *Skp* evolution matched genome evolution, we observed a broad congruence between the two (Supplementary Figure 8), which implies that if recombinatorial exchange occurs as proposed, it most likely occurs among the members of the same population rather than recombining with distant donors. Altogether, these results suggest that recombination can indeed account for how staggered, lineage-specific boundaries average into a continuous valley rather than discrete per-genome cliffs. However, because the recombination signal is genome-wide, the answer to why the sequence divergence is concentrated at *Skp* must therefore be sought elsewhere.

Another potential mechanism worth considering is epistatic co-selection. Recent studies show that genes in close genomic proximity experience markedly elevated rates of co-selection (Cummins et al. 2026), and sequence variation within and between neighboring operon genes can be maintained by shared selective pressures (Mallawaarachchi et al. 2024). The dN/dS of *Skp* was below unity at 0.56, yet it is an order of magnitude above the dN/dS of the genes that flank this operon (0.06, on average), and far above the genome-wide norm for this lineage, where over 95% of the core genes fall below pN/pS of 0.15 (Kiefl et al. 2023). Against this backdrop of pervasive, intense purifying selection that acts on this clade, the dN/dS of *Skp* suggests a gene-specific relaxation that leaves room for diversifying selection to align *Skp* diversity to its OMP clients. If selection indeed acts primarily on *Skp* through its OMP clients, then the remaining genes in this operon that synthetize and process the very substrates those clients ultimately interact may experience correlated selective pressures that weaken with functional distance from *Skp*. Three lines of evidence support this narrative. First, the relaxation of purifying selection was not abrupt but graded: dN/dS declined steadily from 0.56 at *Skp* through its immediate neighbors (*BamA*, *Dxr*, *CdsA*; 0.25 to 0.39) to 0.06 in the flanking genes, revealing a selective gradient centered on the chaperone (Supplementary Figure 9a). Second, when substitutions in each gene were mapped onto the genome phylogeny, the operon genes co-diverged with *Skp* well beyond the genome-wide null expectation for ‘more time, more change’ (mean rate-controlled partial correlation of 0.54, versus 0.22 for flanking genes; Mann-Whitney p = 3.9e-3). Moreover, pairwise co-divergence among the operon genes formed a coherent block rather than decreasing with genomic distance, suggesting that the genes in the operon share a selective regime rather than diverging independently (Supplementary Figure 9b,c). Third, and most critically, this co-divergence signal extended beyond the operon itself to the functional partners of *Skp* throughout the genome: envelope-biogenesis genes (COG category M) were enriched two-fold among the top decile of *Skp*’s co-divergence partners (15.2% vs 7.1%; hypergeometric p = 3.5e-3) and showed higher partial correlations than the rest of the genome (Mann-Whitney p = 4.7e-8; Supplementary Figure 9e). The strongest partners of *Skp* included the OMP-assembly factor *BamD*, the *LptBFGC* LPS-export system, and the protein-handling chaperones *SecB* and *GrpE*. This enrichment did not appear to be a rate artifact as the envelope genes still exceeded their peers when we ranked them only against substitution-count-matched genes (median 60th percentile; Wilcoxon p = 1.9e-3), just like did *LptD* and *BamA* (80th and 87th percentiles; Supplementary Figure 9f). This rules out the most parsimonious alternative that any fast-evolving gene would appear to co-diverge with *Skp*, and instead shows that *Skp* and the genes involved in envelope biogenesis evolve under shared selective pressures arising from functional coupling, giving further credence to epistatic co-selection as a mechanism to explain the divergence gradient. A striking example of this phenomenon was the phylogenetic congruence between *Skp* and *SurA*, another periplasmic chaperone that works synergistically with *Skp* to prevent OMPs from misfolding during their passage through the periplasm (Chamachi et al. 2022). Even though *SurA* was encoded in region #226, which was over 195,000 nucleotides away from *Skp* in every genome, with its own strong divergence gradient, the evolution of *SurA* tracked that of *Skp* more closely than either of these genes tracked the ancestral relationships between genomes (Mantel r = 0.65, p < 1e-4) (Figure 6).

**Figure 6:**
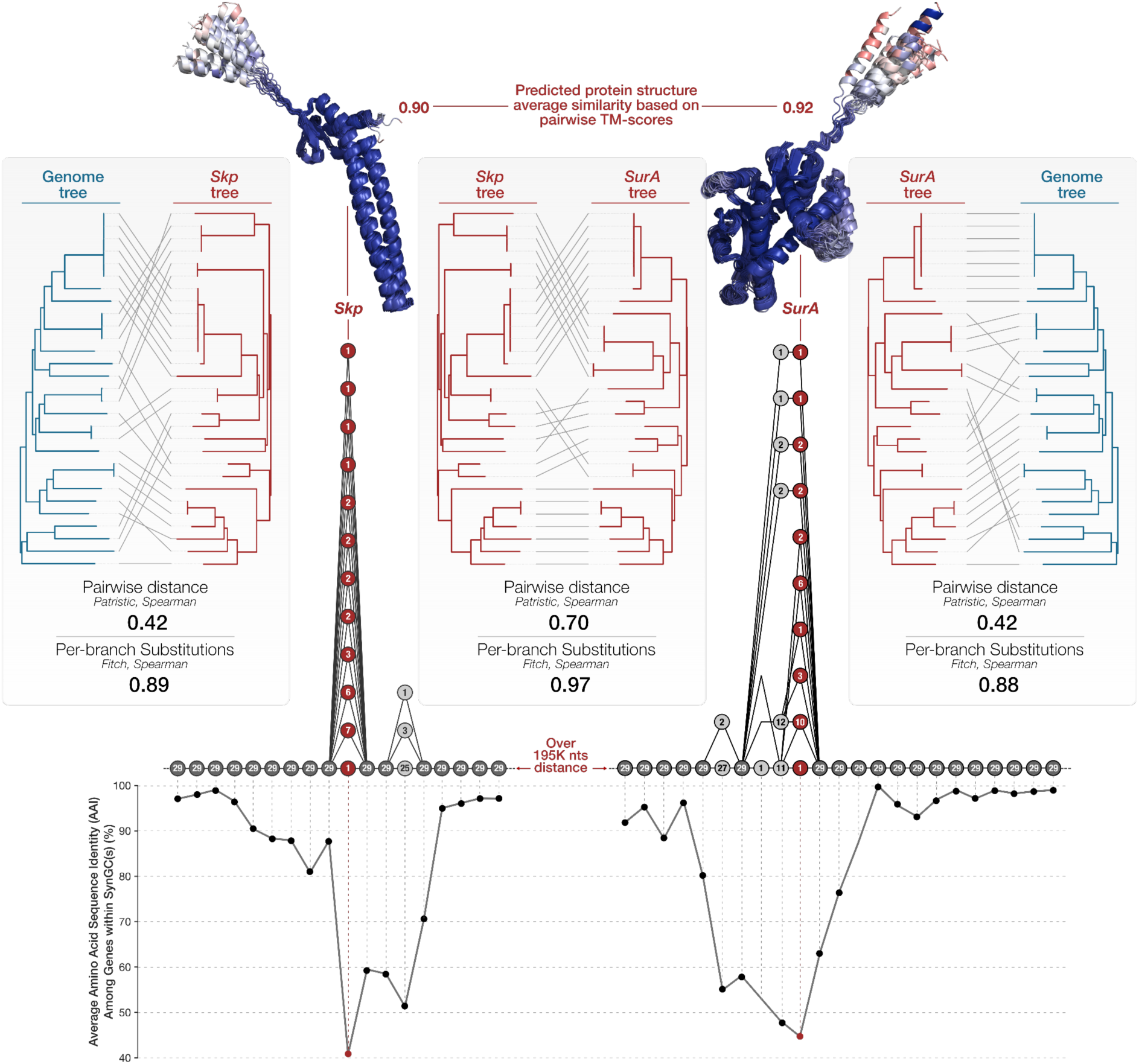
Comparison of *Skp* vs *SurA* vs genome trees. Pairwise tanglegrams display three different comparisons: from left-to-right, (1) the genome tree versus the *Skp* tree, (2) the *Skp* tree versus the *SurA* tree, and (3) *SurA* tree versus the genome tree. Under each tanglegram, the congruence between the two trees are shown as the patristic-distance (Mantel) and per-branch (Fitch) Spearman correlations. The figure also includes the pangenome subgraphs for the region that encodes *Skp* and the one that encodes *SurA* along with within-SynGC average amino acid sequence identity of all genes. The superimposed protein structures show the congruence between AlphaFold2-predicted protein structures for all *Skp* sequences and all *SurA* sequences. The structures are colored based on structure homology calculated with US-align.

These observations suggest that the sequence similarity landscape around *Skp* yields a multifaceted story: pervasive recombination reshuffles and redistributes divergent alleles among genomes, whereas epistatic co-selection shapes and localizes that divergence, concentrating it on *Skp* and grading it outward through the operon. Our data also show that the co-divergence signal is not limited to genomic neighbors but extends to envelope biogenesis genes dispersed across the *Undatipelagibacter* chromosome, including *SurA*, a functional partner of *Skp* (Figure 6, Supplementary Figure 9e, 9f), suggesting that proximity-based co-selection is only one facet of a broader evolutionary process in which an entire functional subsystem is maintained as a single evolutionary unit, with co-varying genes regardless of genome location. Insights into the precise evolutionary and molecular mechanism that maintains such patterns and their functional targets require a larger number of genomes across the tree of life. But our ability to detect and quantify such fine-grained, position-specific divergence gradients within otherwise conserved operons demonstrates the analytical power of the synteny-aware pangenome graphs to reveal a landscape of variation that is typically lost in conventional pangenomes.

## Conclusions

Conventional pangenomics has transformed comparative genomics by quantifying gene content variation across closely related organisms, yet by discarding synteny it also obscured the rich architectural organization of microbial genomes. By resolving multi-copy gene ambiguities, converting complex graphs into directed acyclic structures, and rendering them through an interactive layout that preserves chromosomal context, our synteny-aware, gene-centric pangenome graph approach makes the full spectrum of genomic variation, spanning large, functionally specialized hypervariable islands to subtle, single-gene divergence gradients, accessible to exploration and quantification.

*Undatipelagibacter* pangenome graph reveals that genomic variability is not a binary property of hypervariable genomic islands and static backbones, but a continuum in which variable regions differ from one another in scale, structural topology, functional identity, and evolutionary character. The composite variability score we developed to capture this continuum, and the structural clustering it enables, provides a quantitative vocabulary for describing and comparing patterns of variation that were previously characterized only qualitatively or remained completely invisible.

The resolution provided by gene-centric synteny-aware pangenome graphs also has direct implications for theoretical models of pangenome evolution. Frameworks that explain the maintenance of distributed gene pools typically treat genes as independent units with individual costs and benefits. Our findings show that genomic variability is organized into functionally coherent, structurally distinct regions that are in some cases evolutionarily ancient and persist across divergent clades, and other cases respond to co-selective pressures through epistasis. These observations suggest that the effective number of independently varying genes in a pangenome may be substantially smaller than naive gene-level analyses imply, and that the evolutionary forces that maintain these units may remain incomplete without the inclusion of their chromosomal context.

As high-quality genomes continue to accumulate across the tree of life, we anticipate that synteny-aware pangenome graphs will serve as a foundation for systematic comparisons of genomic architecture within and between taxa, connecting the patterns of variation visible in graphs to the ecological and evolutionary forces that shape them.

## Materials and Methods

### Synteny-aware pangenome graph computation

Our approaches to compute and visualize pangenome graphs, which we detail below, are captured by two new programs, ‘anvi-pan-genome-graph’ and ‘anvi-display-pan-graph’, that we have implemented in the open-source software ecosystem anvi’o (Eren et al. 2015, 2021). The URL http://merenlab.org/pangraph-tutorial/ serves a step-by-step user tutorial on their application starting from FASTA files.

To reconstruct directed acyclic pangenome graphs and to visualize them interactively, we developed three interconnected algorithms that address (1) synteny-aware gene cluster construction, (2) directed acyclic graph conversion, and (3) determination of the final graph layout. The following subsections describe the details of each step, which start with a list of conventional gene clusters (GCs) that describe homologous genes across a set of genomes, and yield a topologically ordered representation of their genomic context.

### Synteny-aware gene cluster construction

Gene clusters (GCs) in conventional pangenomes can contain multi-copy genes, which introduce circularity when GCs are naively connected in a graph as nodes where synteny between each gene cluster are represented by edges. To resolve this critical complexity issue caused by multi-copy genes, we decompose each GC into one or more ‘synteny-aware gene clusters’ (SynGCs), where each SynGC contains at most one gene per genome, and groups only those multi-copy genes that share the same conserved syntenic context (where any SynGC describes the only set of homologous genes that share the exact syntenic context). This outcome is ensured through two complementary comparisons that assess the synteny conservation across all member genes in each GC: First, we evaluate local synteny by comparing the exact gene order in a small neighborhood around each gene copy. Second, we evaluate global synteny by comparing gene content (presence/absence, irrespective of order) across a larger surrounding region. Whether homologous genes from different genomes are placed in the same or separate SynGCs are determined as follows: by default, the local synteny compares ‘gene-mer’ windows that iteratively expand from a minimum of one flanking gene each side until a given gene-mer identifies a unique locus in each genome, while the global synteny examines up to 50 flanking single-copy core genes with a similarity cutoff of 0.5 by default. The user can tighten these thresholds through the command line interface while working with closely related genomes if biologically meaningful syntenic variants are incorrectly merged into the same SynGC, or relax them if SynGCs are over-split and result in highly fragmented graphs for relatively highly divergent genomes. The combination of the two approaches resolves different rearrangement scenarios, such as operon rearrangements that typically have high local but low global synteny, or variable region INDELs that typically have low local but high global synteny. Similar to ‘core’ and ‘accessory’ GCs in conventional pangenomes, we classify the resulting SynGCs in pangenome graphs into multiple categories that consider both their distribution patterns across genomes and their synteny properties (Figure 7): ‘Core SynGCs’, which derive from single-copy core GCs with conserved synteny (where each parent GC yields exactly one core SynGC containing a gene from every genome), ‘Duplication SynGCs’, which derive from multi-copy core GCs with conserved synteny (where each parent GC yields multiple duplicated SynGCs, each containing at most one gene per genome, collectively spanning all gene copies), ‘rearrangement SynGCs’, which derive from core and accessory GCs whose gene copies lack synteny conservation despite presence across genomes, and ‘accessory SynGCs’ or ‘singleton SynGCs’ that derive from accessory or singleton GCs, and appear as such in the final pangenome graph, respectively. Core and duplicated SynGCs constitute the graph ‘backbone’ as they represent SynGCs that describe conserved genes with conserved synteny across all genomes. All other SynGC types populate variable regions, which are always flanked by backbone stretches. In addition to these main categories of SynGCs, the final collection also includes ‘tRNA/rRNA SynGCs’. While these SynGCs do not have a corresponding GC in conventional pangenomes due to their non-coding nature, we include them in the pangenome graph in correct syntenic context due to their relevance to genomic rearrangement events and presence around variable genomic regions.

**Figure 7:**
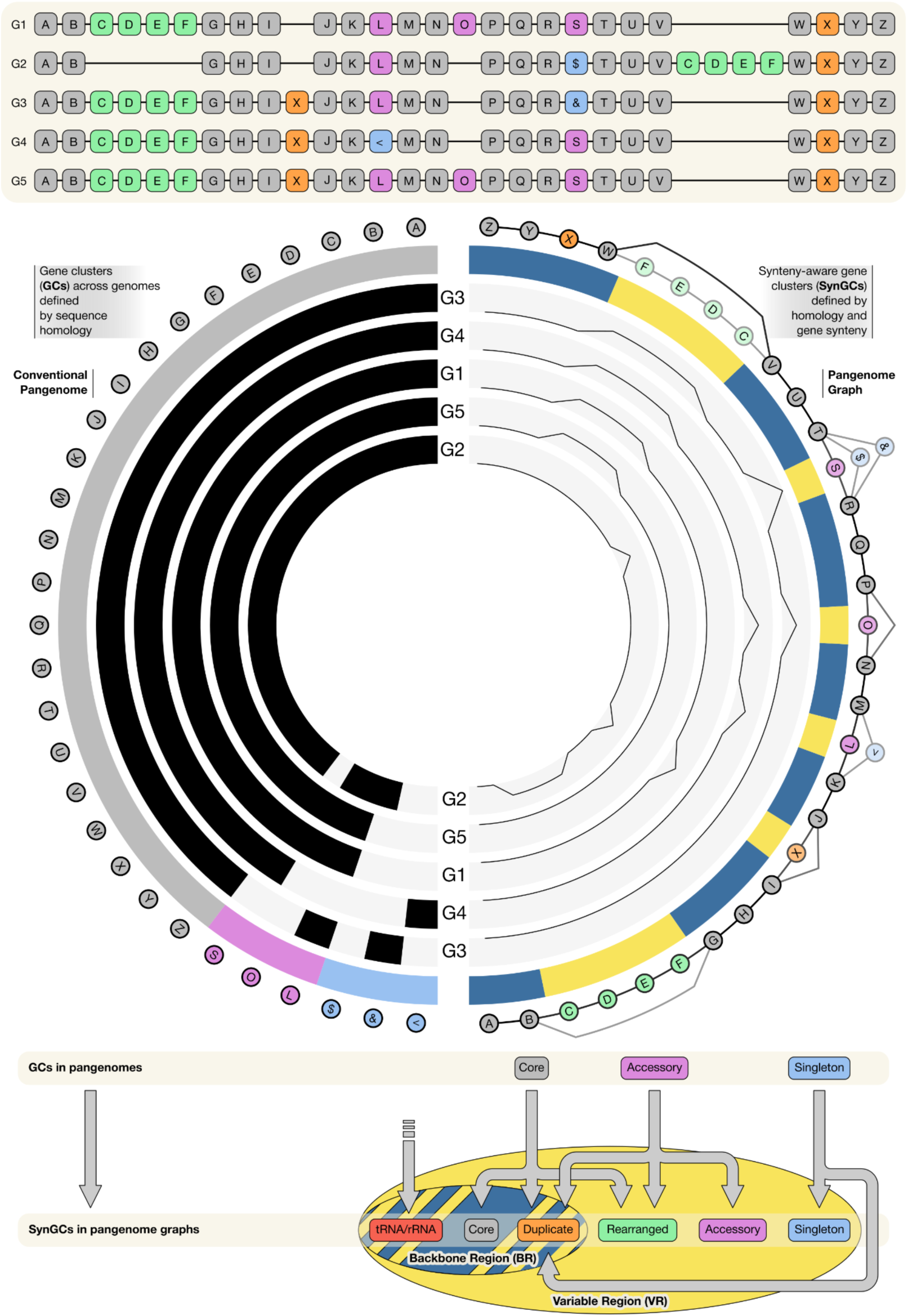
Conceptual pangenome graph workflow. The upper panel shows five example genomes. These genomes contain a rearrangement marked in green, a gene duplication event in orange, single copy core genes conserved in function and position in grey, accessory genes in pink and singletons in blue. The middle panel describes a pangenome created from these five example genomes on the left side and the associated pangenome graph on the right, with the same color coding used in the genomes to describe the present genomic variation. The lower panel of the figure describes how gene clusters (GCs) in the pangenome translate to synteny-aware gene clusters (SynGCs).

### Directed acyclic graph conversion

Even though the decomposition of SynGCs resolves most sources of circularity, residual cycles are often present in the graph as a function of the complexity of the collection of genomes that may include one or more short-range rearrangements with high local and global synteny. We therefore apply a second algorithm to resolve such cases and convert any graph that includes circular features after synteny-aware gene cluster construction into a directed acyclic graph (DAG) by reversing a minimal set of directed edges. Our algorithm first computes a maximum arborescence (Edmonds 1967) of the cyclic graph, which finds a directed spanning tree with the maximum total edge weight in a weighted directed graph (Gabow et al. 1986). In simpler terms, this step selects a single best incoming edge for every node in the graph so that all nodes remain connected without forming loops, while maximizing the total confidence of the selected edges. We then identify the longest path in this tree and designate its terminal node as the starting node. Beginning from this artificial start node, our algorithm backtracks through the tree, trying to connect the current node to an already-visited node at each step. At branching points, the algorithm recurses into the longest available subtree first. When a connection to a visited node is found, the edge is then directed from the current node to the visited node. If this opposes the original edge direction, the edge is marked as reversed. Nodes that cannot be connected to visited nodes are left open and revisited in subsequent passes from the tree root. The procedure terminates when all nodes are connected, yielding a DAG with a minimal set of artificially reversed edges. If this procedure fails and achieving a DAG is impossible, which can happen as a function of the complexity of the input genome or the phylogenetic distance between them, the algorithm terminates.

### Graph layout determination

Having a directed acyclic graph is an important step for the downstream graph visualization and analysis steps, but not sufficient. To visualize the graph that is compatible with genome topologies, we developed a custom layout algorithm to represent the graph in a custom-built interactive interface in anvi’o (https://anvio.org). Our network layout sorts nodes topologically and assigns appropriate x-coordinates to establish a left-to-right ordering of the genomic context. Because the DAG conversion guarantees that every branch eventually merges back into the main path, the algorithm identifies all branching regions, measures their length and nesting complexity, and sorts them from shortest to longest. Y-coordinates are then assigned iteratively: shorter branches are placed closer to the main path, with longer branches layered outward. This inside-out ordering minimizes edge crossings in the final visualization (Figure 7). The interactive interface automatically assigns different colors for the backbone and variable regions on the graph, as well as automatically coloring nodes to distinguish different SynGC categories from one another. However, the interactive interface allows the user to change the visual style and store their preferences.

### Quantifying subgraph topologies and the calculation of the Composite Variability Score

We employed four different measurements to quantitatively describe a given variable region (VR). Each metric explains a different topological aspect of the variable region. The additional terms used within the metrics calculations are listed below.

Let G = {*g*_1_, *g*_2_, … , *g_G_*} be the set of genomes in the dataset and *G* = |G| its cardinality.

Let ℍ = {ℎ_1_, ℎ_2_, … , ℎ*_H_*} be the set of genomes in which the VR is present and *H* = |ℍ| its cardinality.

Let K = {*k*_1_, *k*_2_, … , *k_K_*} be the set of distinct SynGCs in the VR and *K* = |K| its cardinality.

Let ℙ = {*p*_1_, *p*_2_, … , *p_P_*} be the set of unique synteny pathways in the VR, ordered such that:

|*f*(*p*_1_)| ≤ |*f*(*p*_2_)| ≤ ⋯ ≤ |*f*(*p_P_*)| where *f*(*p_i_*) ⊆ G is the set of genomes in which pathway *p_i_* occurs and *P* = |ℙ| its cardinality.

Let *e_i_* = |ℎ(*g_i_*)| be the number of genes contributed by genome *g_i_* ∈ G.

Let *n_i_* = |*t*(*k_i_*)| be the number of supporting genomes *t*(*k_i_*) ⊆ G containing SynGC *k_i_* ∈ K.

**Complexity (*C*)** answers “How many distinct structural realizations (“paths”) occur from the supporting genome?” by estimating the number of events leading to the degree of variation visible in the region. Unique pathways inside the VR are visited in order by the number of genomes backing it, starting with the ones that are less well represented within the dataset. Every visit of a pathway that includes at least one genome not already seen before with this method, counts as one additional degree of complexity, until the breadth of genomic contribution includes every genome of the dataset. Afterwards we subtract one from the result to set backbone regions with just a single straight pathway to zero. The result Y is decreased by one to compensate for the fact that a single pathway is always present and then divided by the number of genomes in the dataset, to weight how high the region can score. Fewer genomes result in fewer possible pathways.

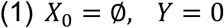

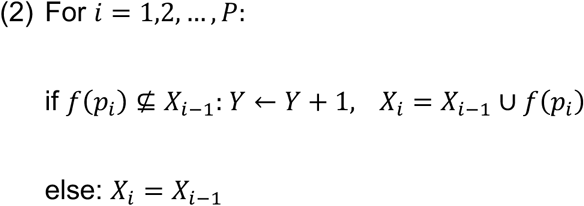

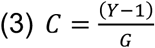

**Expansion (*E*)** answers “*How much gene content is inserted in this variable region?*” by calculating the maximum number of newly introduced genes by a single genome in the region.

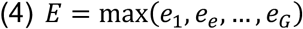

**Diversity (*D*)** answers “*How evenly is genome membership distributed across SynGCs in a region?*” by summarizing how uniform SynGC prevalence is within a region. It first calculates the prevalence proportion *m_i_* of every SynGC in the region, which is the fraction of the genomes in the dataset that contribute to the SynGC and the genomes that contribute to the overall region, and take the variance of these proportions across all SynGCs. The variation is smallest when all SynGCs occur at similar prevalence, and it is largest when they split into rare and ubiquitous extremes. We report the final estimate in reversed form by subtracting the actual variance from the maximum variance possible: the variance of a hypothetical region composed only of conserved SynGCs that are unique to each genome present in the region. Because of this reversal, *D* is the highest when genomes that contribute to a given region are evenly split across all SynGCs in the region.

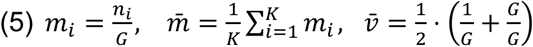

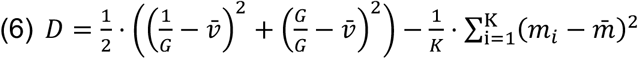

**Weight (*W*)** answers **“***How high is the variable region’s impact?*” by describing the potential significance of a given VR within the genomic landscape through the fraction of *H* and *G*.

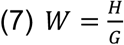

**Composite Variability Score (CVS)** calculates the degree of genomic variation inside a given VR. We use the geometric mean to balance four different terms, requiring higher scores in all metrics to reach a high CVS score.

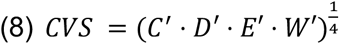

For the calculation of the CVS, all terms are normalized according to min-max normalization

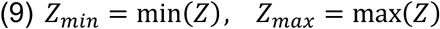

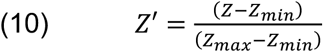

### Generating the *Undatipelagibacter* pangenome

We downloaded the 29 circular genomes for *Undatipelagibacter* as FASTA files from the NCBI (PRJNA1170004). Before the processing of the FASTA files, we ‘reoriented’ them with respect to one another using the program ‘anvi-reorient-genomes’ (https://anvio.org/m/anvi-reorient-genomes), which we implemented for this work to support pangenome graph calculations in anvi’o, and ran it with the flag ‘--use-dnaa-for-reference-orientation’ to ensure that all genomes were aligned and the first gene in the rotated sequences was the *DnaA* gene. We then used these FASTA files to start the anvi’o pangenomics workflow using the program ‘anvi-run-workflow’ (Shaiber et al. 2020), which automates various bioinformatics tasks using Snakemake (Mölder et al. 2021), including gene calling, functional annotation, pangenome creation and ANI calculation. We configured the pangenomics workflow to (1) convert the input FASTA files into anvi’o contigs-db artifacts using the program ‘anvi-gen-contigs-database’ (https://anvio.org/m/anvi-gen-contigs-database), during which Prodigal v2.6.3 (Hyatt et al. 2010), via the Python binding Pyrodigal v3.7.1 (Larralde 2022), called the open reading frames (ORFs), (2) run ‘anvi-run-ncbi-cogs’ to annotate genes in contigs-db files with functions defined in the NCBI’s Clusters of Orthologous Groups (COGs) (Galperin et al. 2025) database through DIAMOND v2.1.12 (Buchfink et al. 2021), and (3) ran the programs ‘anvi-gen-genomes-storage’ (http://anvio.org/m/anvi-gen-genomes-storage) to generate an anvi’o genomes-storage artifact and ‘anvi-pan-genome’ (https://anvio.org/m/anvi-pan-genome) to generate an anvi’o pan-db artifact, which stored the results for the final pangenome for all input genomes as described before (Delmont and Eren 2018). The anvi’o genome-storage and pan-db artifacts generated by the pangenomics workflow represent the primary input files for the pangenome graph calculation.

### Generating the *Undatipelagibacter* pangenome graph

We used the program ‘anvi-pan-genome-graph’ to calculate the pangenome graph with default parameters, and the program ‘anvi-display-pan-graph’ to interactively visualize it. The computation of the pangenome graph for *Undatipelagibacter* from an existing anvi’o pangenome took seven minutes on a high-end laptop computer, all network structures were calculated via NetworkX v3.1 (Hagberg et al. 2008). We summarized the pangenome graph into flat text files for downstream analyses using the program ‘anvi-summarize’.

### Calculating pairwise sequence identity values

We used the program ‘anvi-compute-genome-similarity’ to calculate the average nucleotide identity between pairs of genomes, which relied upon pyANI v0.2.12 (Pritchard et al. 2016). To calculate pairwise identities of arbitrary sequences we used the Python library ‘biopython’ v1.86 (Cock et al. 2009).

### Statistical tests

To test whether there was a linear relationship between complexity and expansion estimates for VRs generated by the program ‘anvi-pan-genome-graph’, we calculated the Pearson Correlation Coefficient (r) using the Python library ‘scipy’ v1.15.2 (Virtanen et al. 2020). To test whether CVS values across variable regions form a continuous distribution or fall into distinct groups (e.g., “hypervariable” versus “other”), we applied three complementary statistical tests to the CVS values reported by the program ‘anvi-pan-genome-graph’. First, we used Hartigan’s dip test (Hartigan and Hartigan 1985) to assess unimodality of the CVS distribution. Second, we applied the Pruned Exact Linear Time (PELT) changepoint detection algorithm (Killick et al. 2012), as implemented in the Python library ‘ruptures’ (Truong et al. 2020), to the rank-ordered CVS values using an L2 cost model and a BIC-derived penalty (ln(n), where n is the number of variable regions) to test for any statistically supported inflection points in the ranked curve. And finally, we compared the fit of a single log-normal distribution (fitted via maximum likelihood) to two-component and three-component Gaussian mixture models fitted using the expectation-maximization algorithm implemented in the Python library ‘scikit-learn’ (Pedregosa et al. 2012) before selecting the best model by the Akaike Information Criterion (AIC) and Bayesian Information Criterion (BIC).

### Protein structure prediction

We used Colabfold v1.5.5 (Mirdita et al. 2022) to predict protein structures from amino acid sequences and US-align v20241201 (Zhang et al. 2022) to create multiple structure alignments and calculate ‘TM-scores’, which represent the length-normalized structural similarity that ranges between 0 and 1, where 0.5 or above indicates similar folds (Zhang and Skolnick 2004). We calculated TM-scores through pairwise comparison of each structure and averaged the TM-scores of each pair and across all comparisons to get a final score per function.

### Running PPanGGOLiN

We used PPanGGOLiN v2.2.5 (Gautreau et al. 2020) to qualitatively compare the pangenome graph calculated with it to the one calculated from our tool. For the creation of the pangenome graph, we used the PPanGGOLiN annotation and workflow tool ‘ppanggolin all’ with the 29 isolate genomes. For visualization of the pangenome graph, we used ‘gephi’ v0.10.1 (Bastian et al. 2009), with the Force Atlas 2 algorithm for graph layout computation, with Stronger Gravity activated and a scaling set to 4000, as per recommendation in the PPanGGOLiN documentation.

### Extraction of the *Skp* locus, and tests for recombination and co-selection

To compare divergence, selection, or recombination across the *Skp* locus, we first manually identified the pair of SynGCs on the *Undatipelagibacter* pangenome to identify the extended *Skp*-encoding locus (i.e., the *LpxB*-*UppS* envelope-biogenesis operon together with its flanking genes on either side), and exported the region from every genome using the program anvi-export-pan-subgraph, which provided us with the contiguous locus DNA to perform recombination tests that ignore gene boundaries, and the individual genes with their alignments to perform per-gene divergence and selection tests. We also calculated a reference genome phylogeny for our genomes as the backbone for the co-divergence analyses using the reproducible workflow that was available to us (Freel et al. 2025) which relied upon 165 alphaproteobacterial single-copy core genes and IQ-TREE v3.0.1 (Minh et al. 2020) and used throughout as the backbone for the co-divergence analyses. To profile divergence and selection gene by gene along the locus, we used FAMSA v2.4.1 (Deorowicz et al. 2016) to codon-align gene amino acid sequences so the reading frame is preserved across indels. At each position we summarized divergence as the mean pairwise amino-acid identity over each column. Because a smooth average can hide a mixture of very different pairs, we retained the full per-pair identity distribution and scored its shape with Sarle’s bimodality coefficient with the classical assumption that values above 5/9 cutoff indicate a two-moded distribution. To separate selective from neutral divergence we estimated synonymous vs non-synonymous divergence from the codon alignments using the Nei-Gojobori counting method (Nei and Gojobori 1986), and reported them as mean pairwise proportions (pS, pN), by excluding saturated codon pairs with Jukes-Cantor-corrected dN/dS values. To see whether divergent alleles fall into genome-specific blocks rather than spreading evenly, we scored each genome’s allele against the per-position consensus, and arranged the resulting genome-by-gene matrix along the previously calculated genome phylogeny. With the assumption that the averaged divergence valley across the operon cannot distinguish gradual in-place divergence from staggered homologous recombination, we tested for recombination directly on the contiguous locus, and without any consideration of the gene boundaries. For this, we aligned the contiguous locus DNA of all genomes at the nucleotide level with FAMSA and removed columns with gaps and ambiguous bases to retain informative sites that are biallelic. On these clean columns we ran the four-gamete test to build a site-by-site incompatibility matrix and summarized it with the pairwise homoplasy index (PHI) as previously described (Bruen et al. 2006; doi:10.1534/genetics.105.048975). From the same matrix we derived a per-site conflict profile along the locus and the relationship between incompatibility and the distance separating sites (i.e., the diagnostic signature of recombination), while we used a sliding-window nucleotide-identity profile to investigate the sequence divergence valley also without gene boundaries. To establish if the signal was confined to the *Skp* operon, we also computed PHI separately for the operon itself and for the regions that flanks it. To investigate whether the locus in which *Skp* was encoded was held together by selection focused on *Skp*, we explored co-divergence on the genome phylogeny. For each gene we mapped column by column amino-acid substitutions onto the genome tree by Fitch parsimony to obtain a per-branch substitution vector. We first asked whether *Skp* tracks the genome as a whole through two complementary tests of gene-genome congruence (Leigh et al. 2011): we correlated patristic distances from the *Skp* gene tree and the genome tree (Mantel test, with the *Skp* tip labels permuted), and we correlated per-branch *Skp* substitutions against genome-wide branch lengths. We then defined co-divergence between two genes as the partial Spearman correlation of their per-branch substitution vectors controlling for genome-wide branch length (removing the shared “more time, more change” effect so that what remains is gene-specific). We then computed it between *Skp* and every other locus gene and pairwise among all locus genes. To test functional specificity genome-wide, we computed the same partial co-divergence with *Skp* for every core single-copy gene cluster long enough not to be a fragment and carrying enough inferred substitutions to support a correlation, classified genes as cell-envelope associated genes (if they were resolved to COG category M) or other, and tested envelope enrichment among *Skp*’s partners with a Mann-Whitney U test and a hypergeometric test on the top decile. We confirmed the enrichment is not a rate artifact by ranking the co-divergence estimate for each envelope gene against the non-envelope genes closest to it in substitution count and testing those percentiles against the median with a one-sided Wilcoxon signed-rank test.

We performed all these tests using the full set of 29 genomes, and repeated them after a redundancy analysis to remove flat branches from the genome tree and ensure our results are not driven by redundant sampling of the clade: the results were virtually identical, and significance scores remained significant. Throughout these analyses we used anvi’o outputs with functions for statistical and phylogenetic analyses implemented in libraries NumPy (Harris et al. 2020), SciPy (Virtanen et al. 2020), and DendroPy (Sukumaran and Holder 2010). Our online bioinformatics workflow contains code and descriptions of each step that are fully reproducible starting from the anvi’o pangenome graph.

### Postprocessing of figures

We used either anvi’o or matplotlib v3.10.8 (Hunter 2007) in combination with numpy v1.26.4 (Harris et al. 2020), scipy v1.15.2 (Virtanen et al. 2020), pandas v2.3.3 (The pandas development team 2026; McKinney 2010) and seaborn v0.13.2 (Waskom 2021) to generate the figures in our study and finalized them for publication using Inkscape v1.4.3, an open-source vector graphics editor (https://inkscape.org).

## Data availability

*Undatipelagibacter* genomes used in our study are publicly available via the NCBI BioProject ID PRJNA1170004. The raw data, intermediate data products, as well as a reproducible bioinformatics workflow with ad-hoc Python scripts and their applications are available at https://merenlab.org/data/undatipelagibacter-pangenome-graph.

## Acknowledgements

The authors thank Sophie Griesbach, Sean Crosson, Aretha Feibig, Ralf Rabus, and Andreas Teske (who proposed the term ‘Persistent Variable Regions’), as well as the members of our group (https://merenlab.org/people/) at the Helmholtz Institute for Functional Marine Biodiversity (https://hifmb.de) for helpful discussions. We used the HPC cluster ROSA at the University of Oldenburg for heavy duty computational tasks, which is funded through the German Research Foundation (DFG) Major Research Instrumentation Program (INST 184/225-1 FUGG) and the Ministry of Science and Culture (MWK) of Lower Saxony. We thank Stefan Harfst and members of the Department of Scientific Computing at the University of Oldenburg for their help and patience with us.

## Author contributions

Conceptualization: A.H., A.M.E. Formal Analysis: A.H., A.M.E. Data Curation: S.J.T., K.C.F., M.S.R. Investigation: A.H., A.M.E. Resources: K.C.F., M.S.R., A.M.E. Software: A.H., M.S., F.T., I.V., T.C., A.M.E. Validation: S.J.T., F.T., J.O.M., A.S., M.S.R. Visualization: A.H., M.S., A.M.E. Writing - Original Draft: A.H., A.M.E. Writing - Review & Editing: J.O.M. Project Administration: A.M.E. Supervision: A.M.E. All authors read, commented on, and approved the final manuscript. Additional notes on software contributions: A.H. designed and implemented the core algorithms and wrote the majority of the codebase. A.M.E. contributed substantially to software development outside the core algorithmic components. M.S. contributed to the development of the interactive interface. F.T., I.V., and T.C. contributed to testing, refinement, and/or additional feature development.

## Supplementary Information

### Supplementary Tables

**Supplementary Table 1: General genome information.** a) Dataset ANI. Table with all the calculated ANI values between the datasets genomes. b) Genome Info. Table containing all information of the datasets genomes, including metrics such as completeness, contamination, and size.

**Supplementary Table 2: Summary tables of the pangenome and pangenome graph artifacts.** a) PG to PGG artifact Summary. Table that shows the data used to generate the bottom part of Figure 1. b) PGG artifacts Summary. Table that shows how many genes of the VR and BR artifacts were clustered in how many SynGCs and regions. c) PGG SynGC Summary. Shows how many gene calls and SynGCs are clustered as the different SynGC types. d) PG GC Summary. Shows how many gene calls and GCs are clustered as the different GC types.

**Supplementary Table 3: Classification schemas.** a) GC to SynGC Translation. Describes the various examples of how GCs translate to SynGCs. b) COG24 Groups. Table containing the grouping of the COG24 categories into the five groups used to annotate the regions in Figure 2.

**Supplementary Table 4: Functional distributions of pangenome graph regions.** a) COG24 Groups distribution per region. b) Hellinger distance calculations. c) Null distributions for the violins in Figure 2. d) Cumulative COG24 Groups distribution per region.

**Supplementary Table 5: Comparison of taxa from the *Pelagibacterales order*.** a) COG24 Annotation of Pelagibacter region #26. b) COG24 Annotation of Elongatipelagibacter region #176. c) COG24 Annotation of Fontibacterium region #98. d) COG24 Annotation of Litoralipelagibacter region #64. e) COG24 Annotation of Xanthinipelagibacter region #178. f) COG24 Annotation of Atlantikopelagibacter region #142. g) COG24 Annotation of Coralipelagibacter region #162. h) COG24 Annotation of *Undatipelagibacter* region #32.

**Supplementary Table 6: Output tables from anvi’o pangenome graph.** a) Gene Call to SynGC. A result table from the pangenome graph, maps every gene call to a SynGC. b) SynGC Position. A result table from the pangenome graph, maps every SynGC to a position and region. c) Regions. A result table from the pangenome graph, describes every region in detail, e.g. position, complexity, diversity. d) Gene Call Information. A table from the pangenome, describes every gene call in detail, including COG24 functional annotations.

**Supplementary Table 7: Sequence identity per position and functional annotation.** a) AA sequence similarity per position of backbone region, region #148, region #150, region #224 and region #226. b) Data used for ORF prediction, including gene distance and strand position. c) per position functional annotations of the envelope biogenesis operon, as well as the operon upstream and the operon downstream. d) tm scores of the aligned protein structures trimmed with a confidence cutoff of 70 of position 1401. e) tm scores of the aligned protein structures trimmed with a confidence cutoff of 70 of position 1762 f) tm scores of the aligned untrimmed protein structures of position 1401. g) tm scores of the aligned untrimmed protein structures of position 1762.

### Supplementary Figures

**Supplementary Figure 1:**
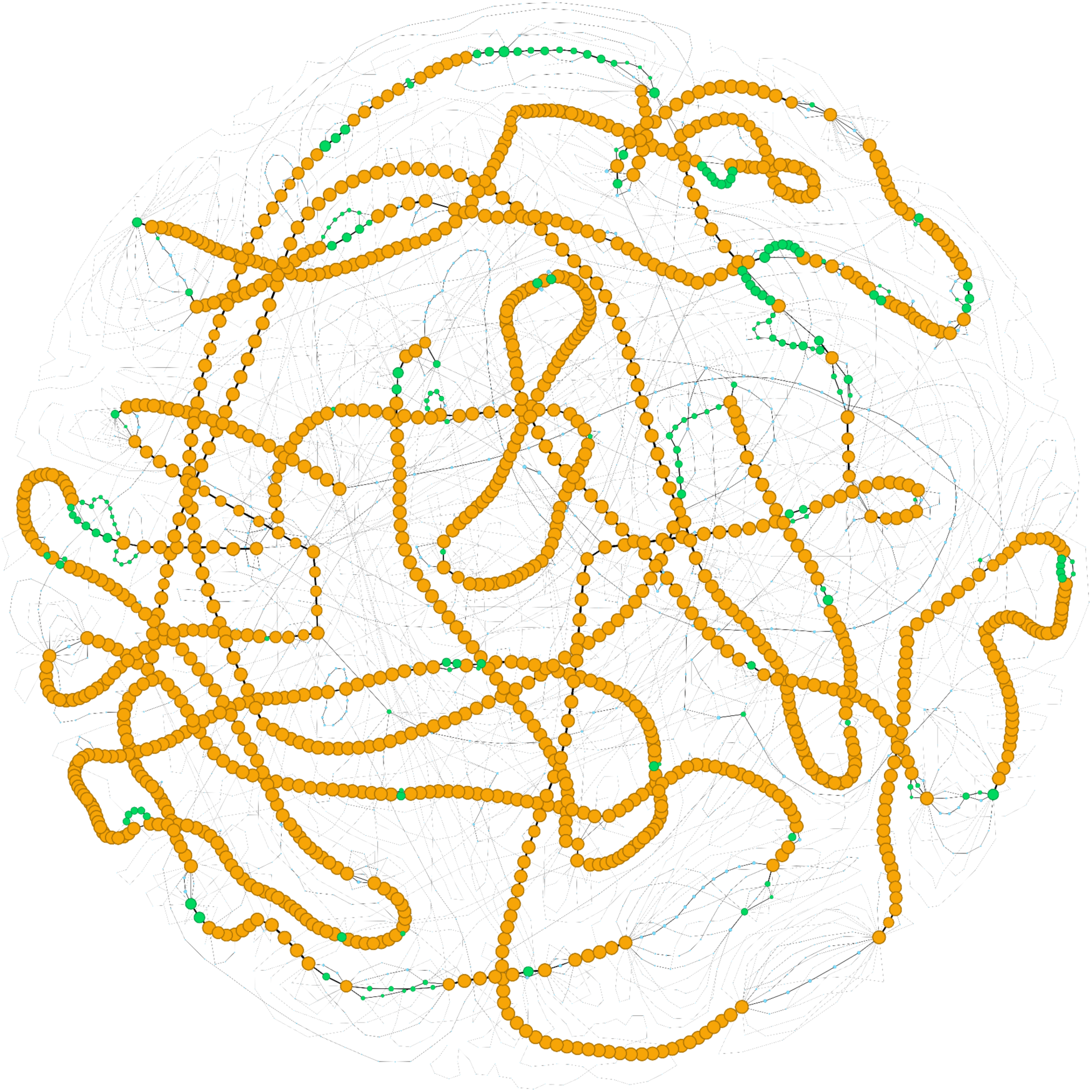
The *Undatipelagibacter* PPanGGOLin pangenome graph. The *Undatipelagibacter* pangenome graph that includes the same 29 genomes we have included in our analyses, as calculated by PPanGGOLiN. Orange nodes indicate PPanGGOLiNs persistent genome, green bullets the shell genome and light-blue bullets the cloud genome. The black lines define gene synteny.

**Supplementary Figure 2:**
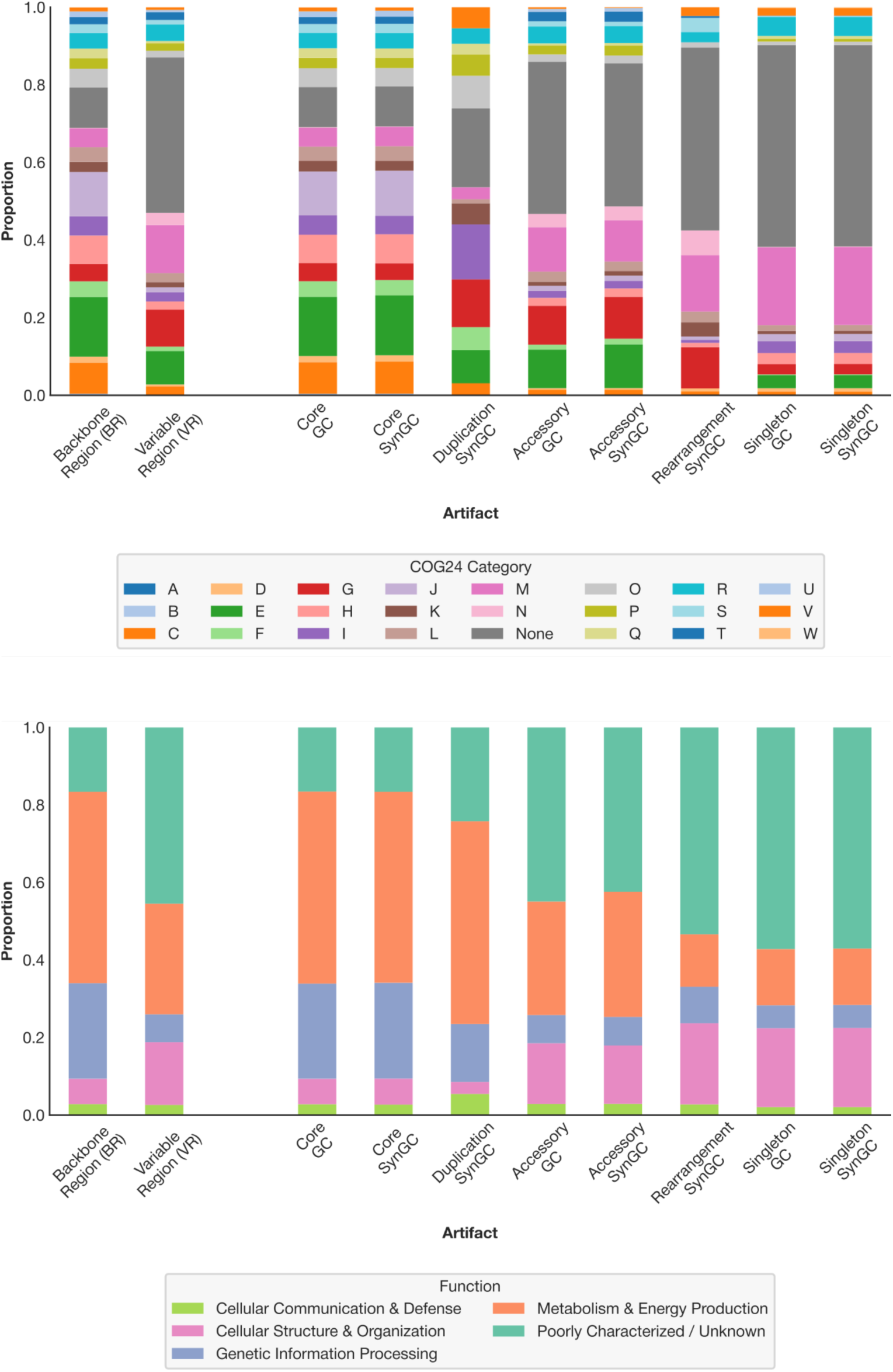
Comparison of Pangenome Graph artifacts. The top panel shows the distribution of COG24 functions between different artifacts of the *Undatipelagibacter* pangenome graph. The bottom panel offers the same information but using the five major groups we used to simplify the COG24 categories instead.

**Supplementary Figure 3:**
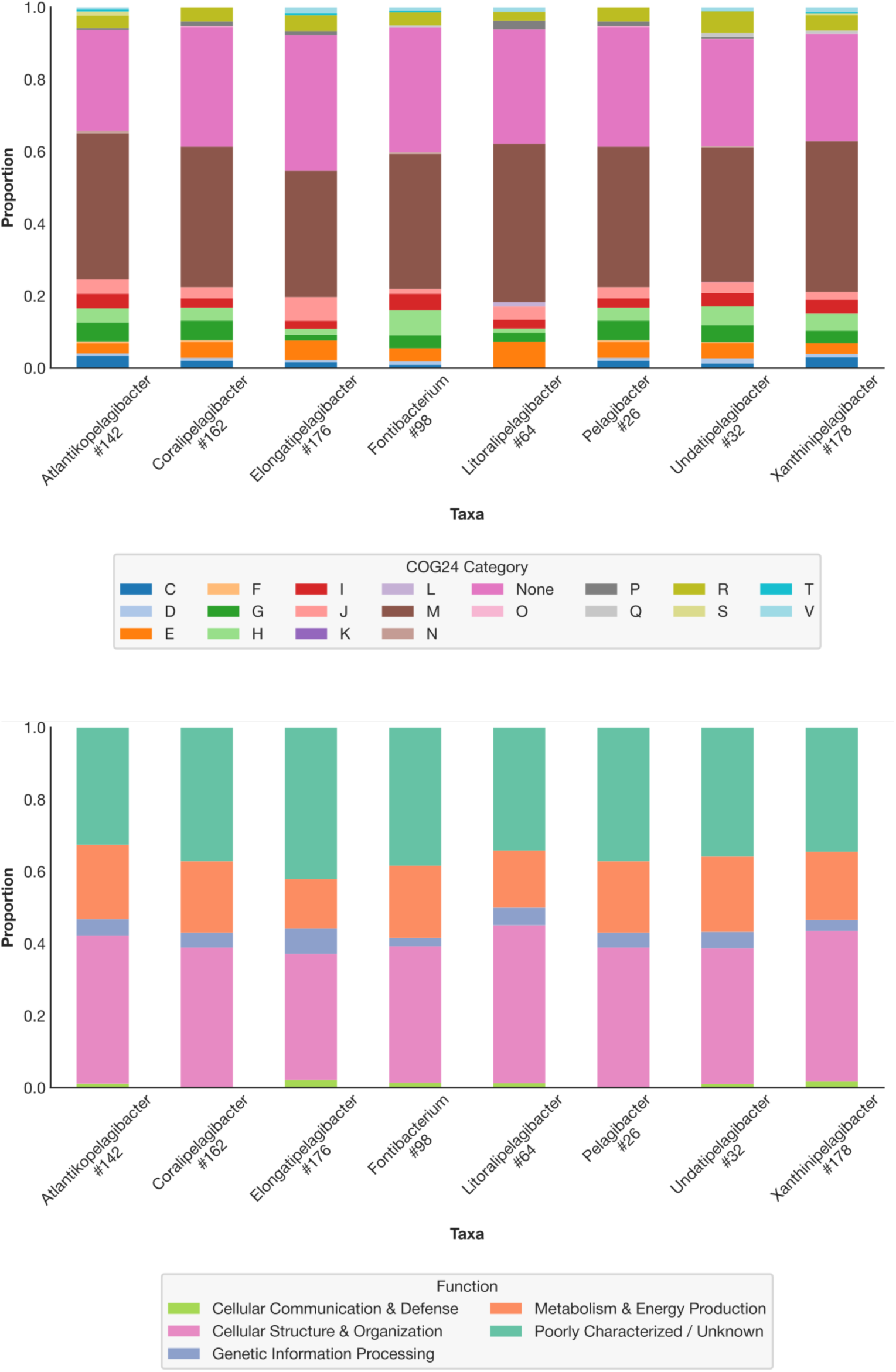
Comparison of pangenome graph regions between different genera of the *Pelagibacterales* order. The top panel shows the distribution of COG24 functions between a similarly located region in pangenome graphs of different genera of the *Pelagibacterales* order. The bottom panel offers the same information but using the five major groups we used to simplify the COG24 categories instead.

**Supplementary Figure 4:**
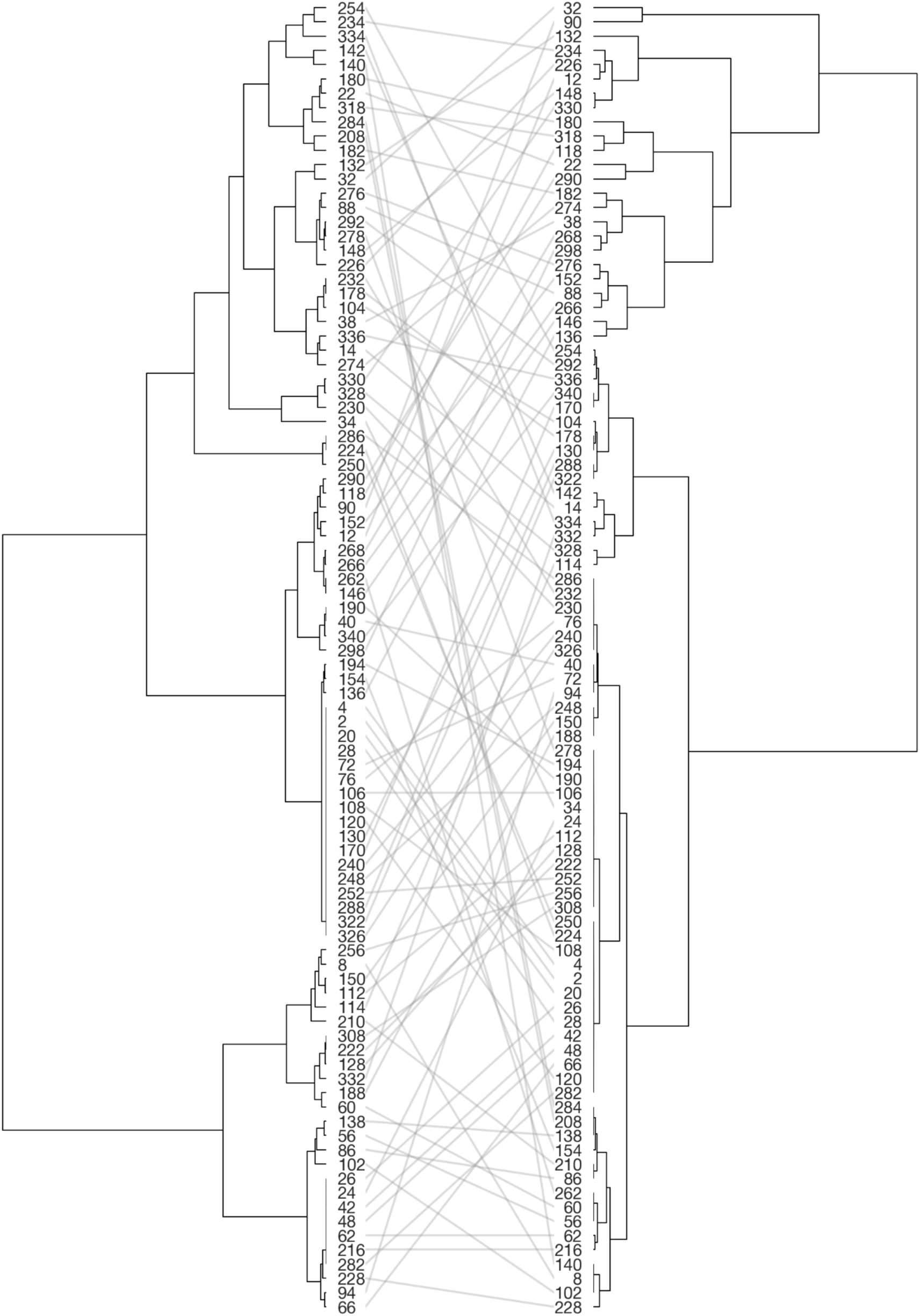
Comparison of the dendrograms created via the functional distribution and metric patterns. The right side is identical to the dendrogram we created with the metrics expansion and complexity for Figure 3. The left side shows the dendrogram we created for Figure 2, with rotated subtrees to minimize the number of line crossings.

**Supplementary Figure 5:**
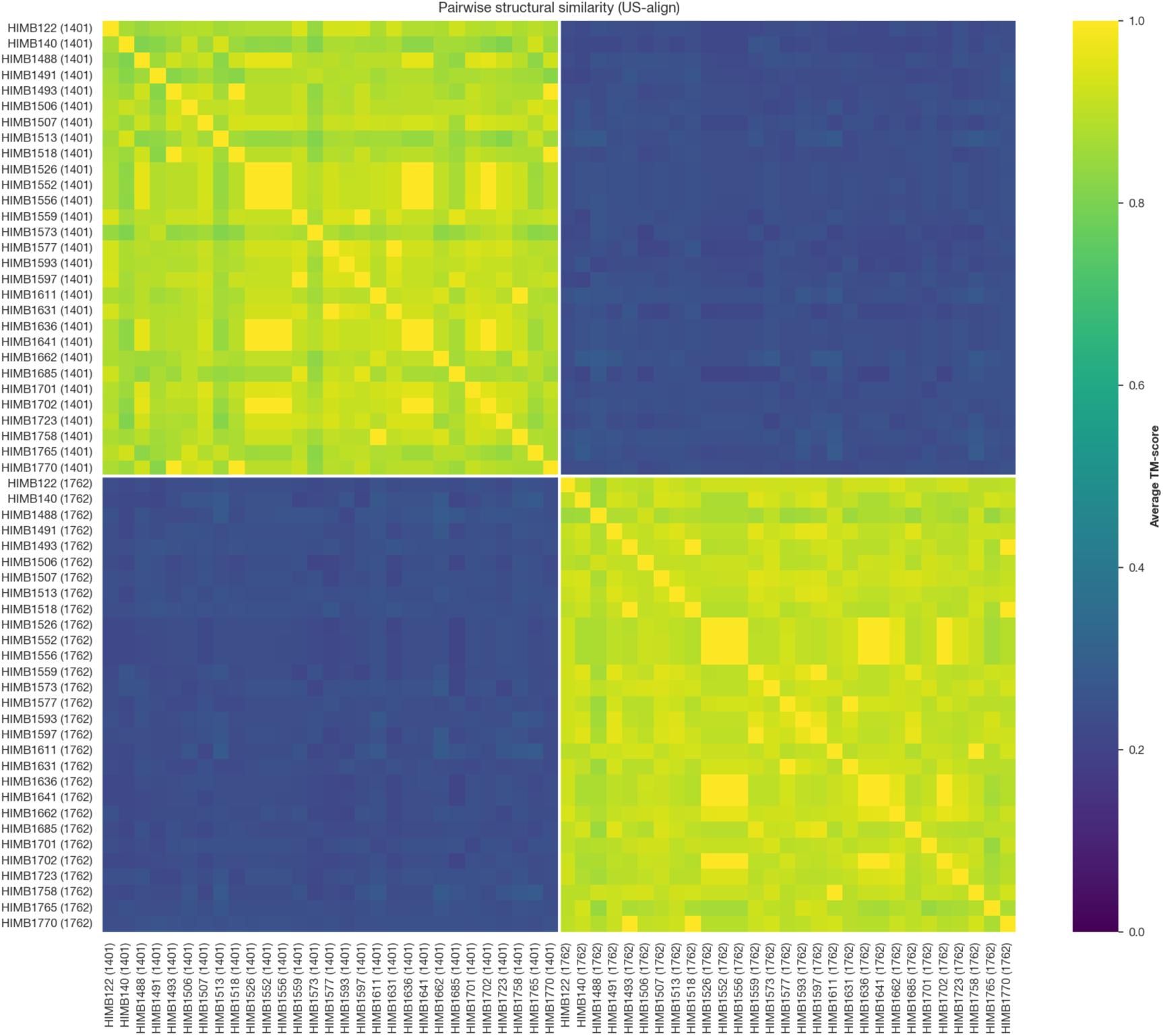
Average TM-scores of Alphafold2 predicted protein structures of *Skp* and *SurA* genes. This figure shows the pairwise structural similarity as the average TM-score of the folded proteins of the genes *Skp* in region #148, position 1401 and *SurA* in region #226, position 1762 in the *Undatipelagibacter* pangenome graph.

**Supplementary Figure 6:**
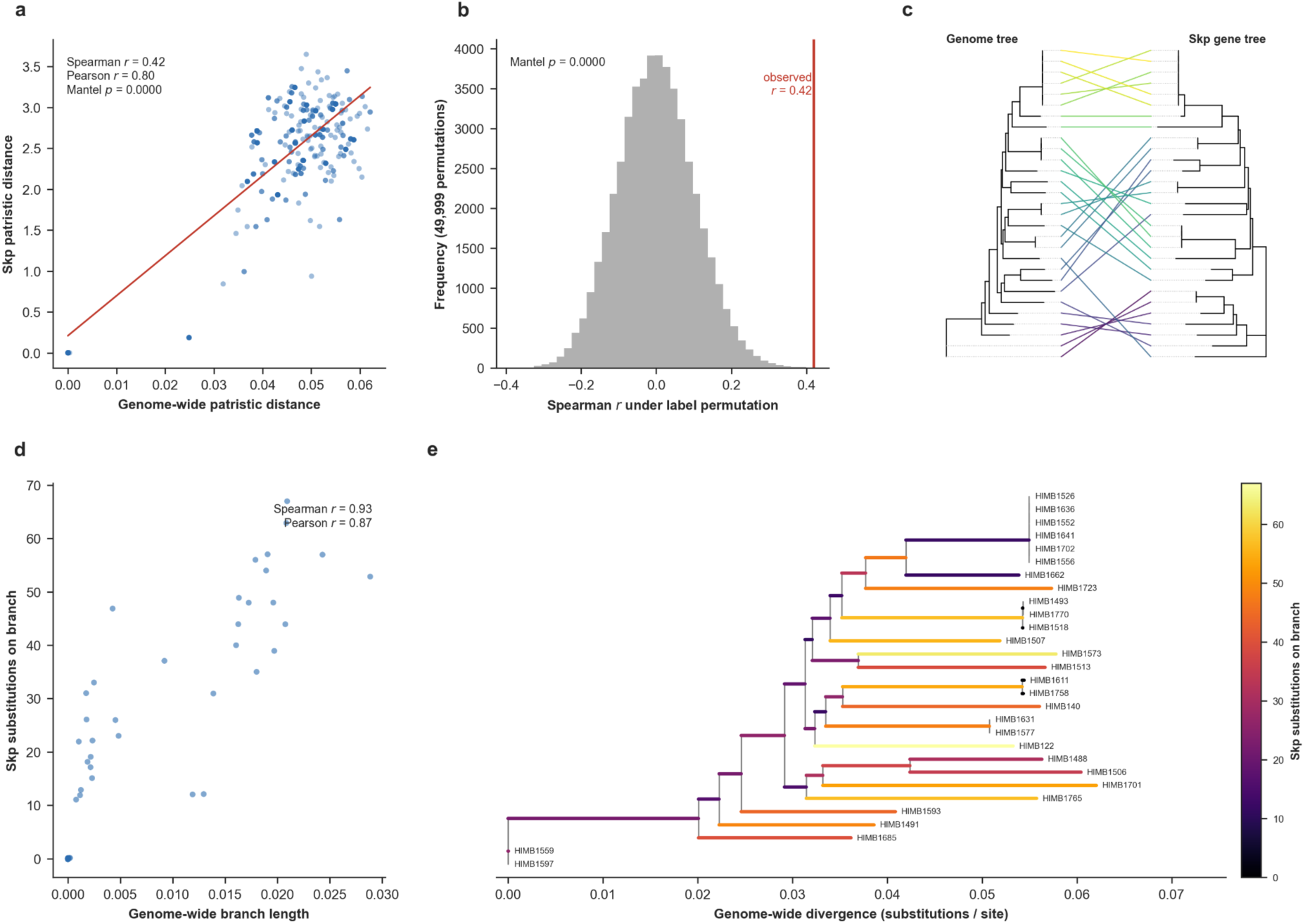
Gene-level signatures of the *Skp* operon sequence divergence gradient and a within-population test for staggered recombination. In every panel the operon that encodes *Skp* is shaded and the *Skp* gene is marked with a dashed line. (a) Mean pairwise amino-acid identity (AAI) at each gene. (b) The same profile drawn for each individual genome pair with the mean overlaid. (c) Distribution of the 406 pairwise AAI values at each gene; red numbers indicate when Sarle’s bimodality coefficient exceeds 5/9. (d) Mean pairwise proportion of nonsynonymous differences per gene (Nei-Gojobori). (e) Heatmap of each genome’s allele identity to the per-gene consensus sequence where genomes are ordered by the genome phylogeny.

**Supplementary Figure 7:**
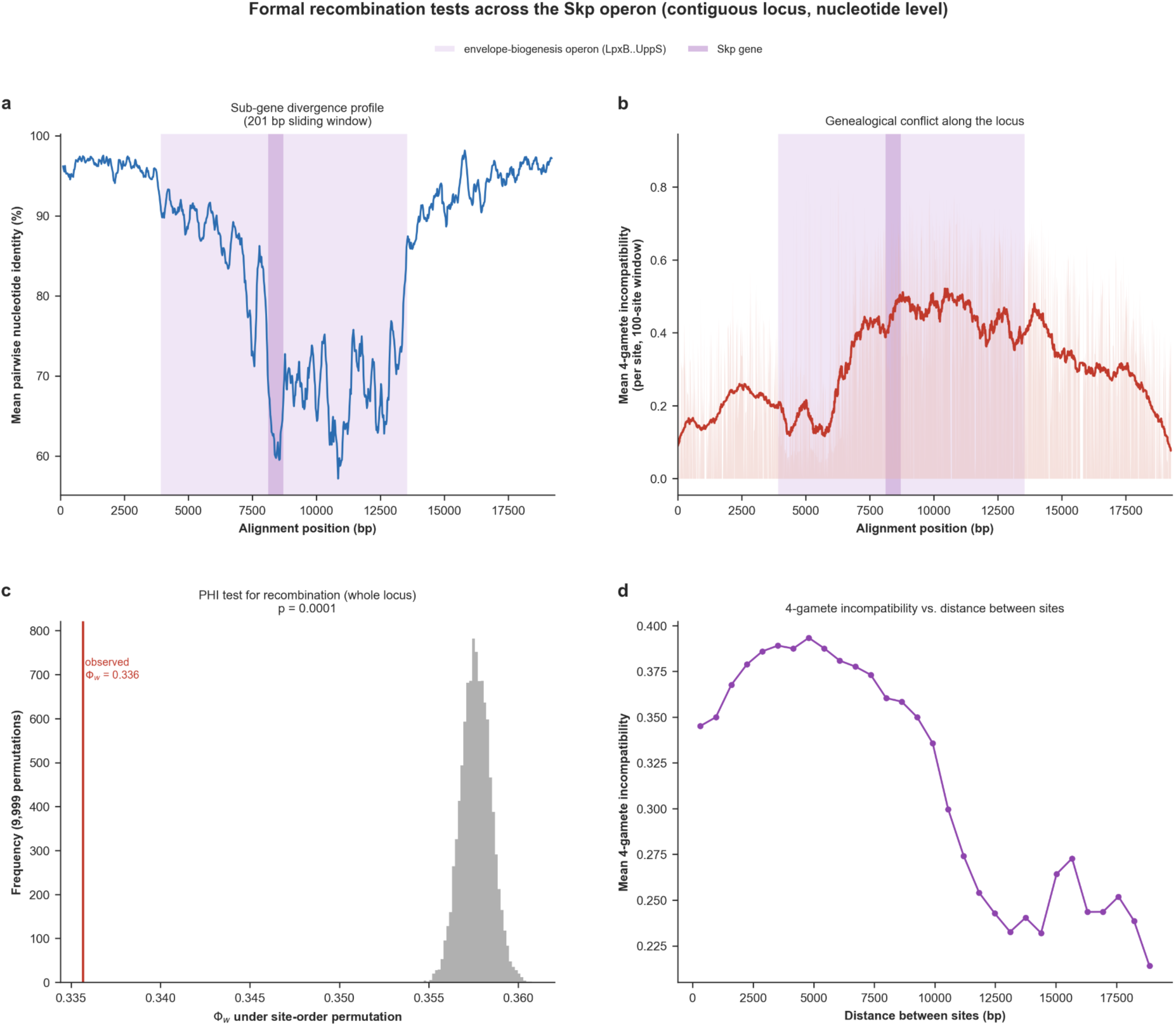
Sub-gene resolution recombination tests across the contiguous *Skp* locus. (a) Mean pairwise nucleotide identity in a 200-bp sliding window along the locus, resolving the divergence valley within and across genes. (b) Genealogical-conflict landscape: for each informative site, the mean 4-gamete incompatibility with its neighboring sites, positioned along the locus. (c) The pairwise homoplasy index (PHI, Φw) test for recombination. The histogram is the null distribution of Φw under 9,999 random permutations of site order; the red line is the observed value. (d) Mean 4-gamete incompatibility between pairs of informative sites as a function of the distance separating them.

**Supplementary Figure 8:**
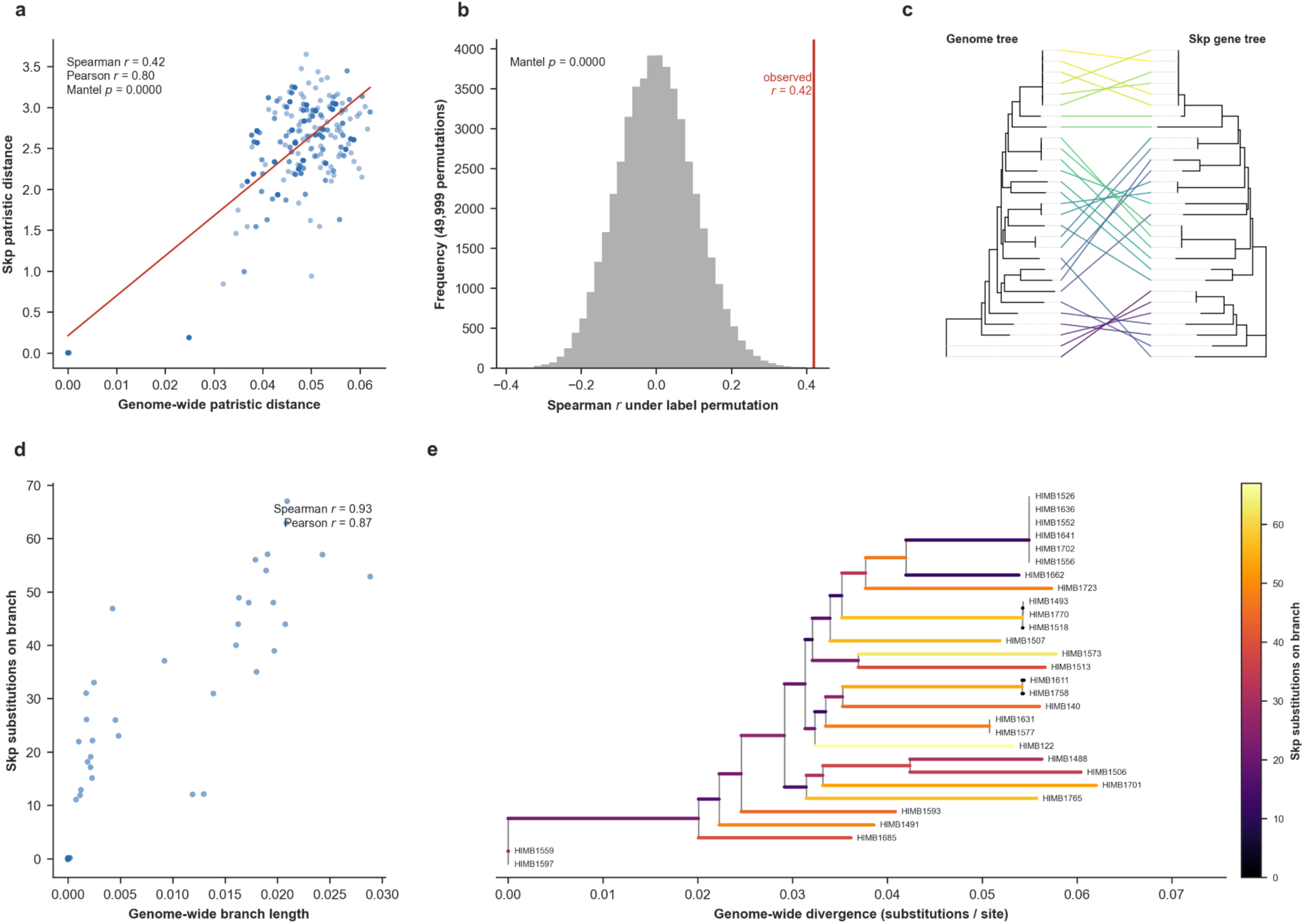
*Skp* divergence versus the genome phylogeny. The top row addresses whether genome-wide and *Skp* pairwise distances agree, and the bottom row addresses whether per-branch *Skp* substitutions scale with genome-wide divergence. (a) Pairwise patristic distance on the genome tree versus on the *Skp* gene tree, with a least-squares guide line. (b) Mantel permutation test. (c) Tanglegram of the genome tree (left) and the *Skp* gene tree (right). (d) *Skp* amino-acid substitutions inferred on each branch of the genome tree by Fitch parsimony, versus that branch’s genome-wide length (substitutions/site). (e) The genome tree with each branch colored by its inferred number of *Skp* substitutions.

**Supplementary Figure 9:**
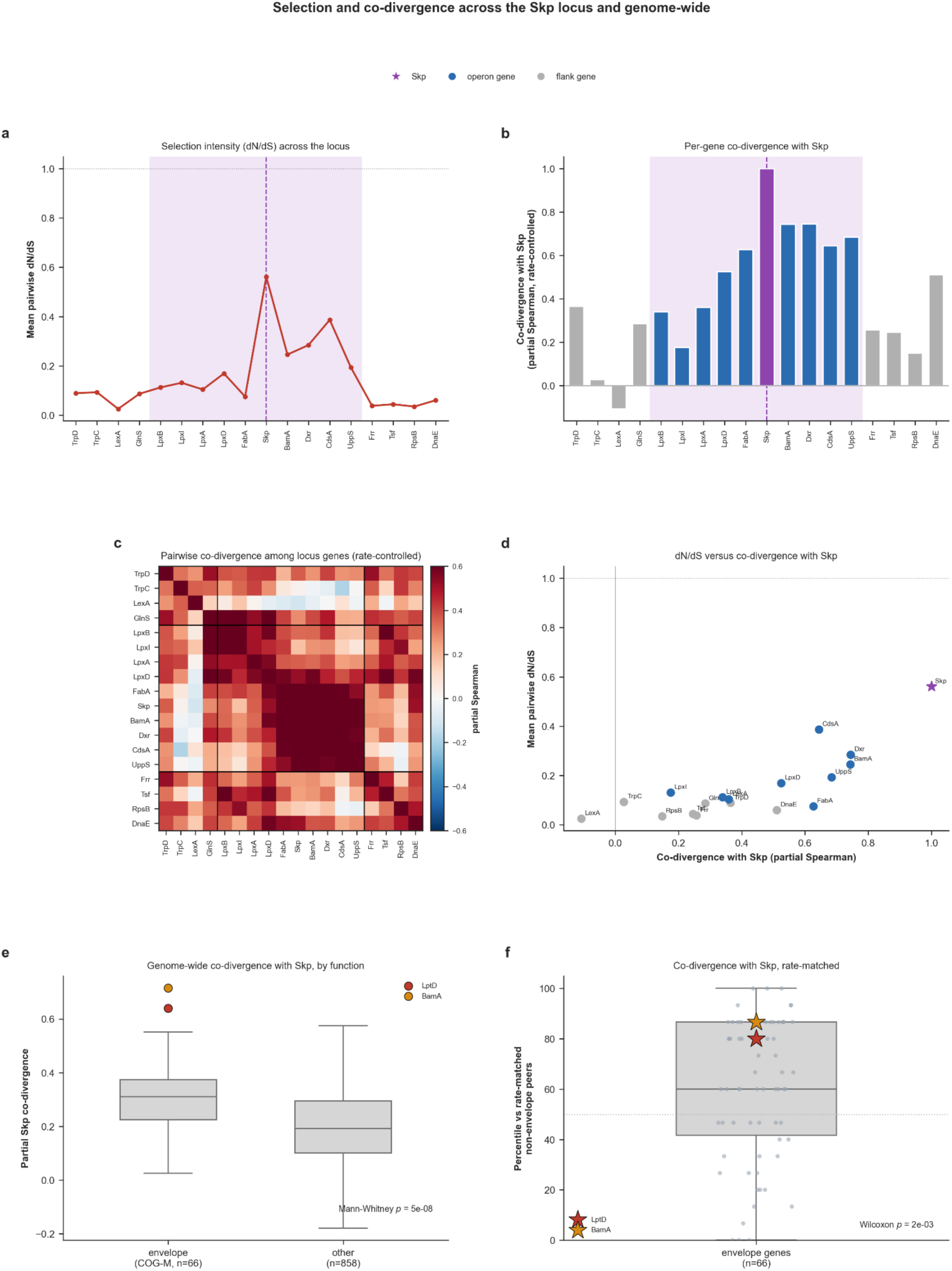
Selection and co-divergence across the *Skp* locus and genome-wide. Operon-internal analyses (a-d) use the per-gene codon alignments of all locus genes with amino-acid substitutions mapped onto each branch of the genome tree by Fitch parsimony, and the genome-wide analyses (e-f) apply the same test to all single-copy core genes. Throughout, ‘co-divergence with *Skp*’ indicates the partial Spearman correlation between the per-branch substitutions of a given gene and *Skp*, while controlling for genome-wide branch length. In a-d the operon is shaded and *Skp* is marked. (a) Mean pairwise dN/dS (Nei-Gojobori) at each locus gene. (b) Each gene’s rate-controlled co-divergence with *Skp*. (c) Pairwise co-divergence among all locus genes. (d) dN/dS versus co-divergence with *Skp*, one point per gene. (e) Genome-wide partial co-divergence with *Skp* for envelope-biogenesis genes (COG category M; n = 66) versus all other single-copy core genes (n = 858), where *LptD* and *BamA*, *Skp*’s most obvious outer-membrane clients, marked. (f) Each envelope gene’s percentile rank for co-divergence with *Skp* among the non-envelope genes closest to it in substitution count.

